# Basal ganglia output guides decision-making by state destabilization

**DOI:** 10.64898/2026.11.01.763532

**Authors:** Yuki Tanimoto, Makio Torigoe, Tanvir Islam, Hisaya Kakinuma, Toshiyuki Shiraki, Merlin Lange, Takuya Isomura, Hitoshi Okamoto

## Abstract

To execute learned behavior, the nervous system can undergo an activity-state transition between the current and the learned activity states. Here, using adult zebrafish, we show that their basal ganglia output induces such state transitions for goal-directed behavior. After learning the Go/No-go task in virtual reality, the striatal indirect pathway responds to negative values of a conditioned cue and the absence of sensory feedback by different neuronal populations. The pallidal output pathway integrates these negative value signals and then broadcasts the integrated signal to the Dc area in the pallium (fish cortex). Optogenetics and mathematical modeling suggested that this pallidal output activity uniformly destabilizes ongoing Dc activity states, thereby facilitating behavioral state transitions toward the behavioral state with the minimum pallidal output activity. This mechanism explains how pallidal output activity guides goal-directed behavior beyond potential barriers between activity states underlying different behavioral states.

## Main text

Animals generate neural ensembles showing attractor-like dynamics to select and sustain specific neural activity states under given conditions.^1–6^ One mechanism for the transition between these activity states is to flatten the potential landscapes of activity states and reduce potential barriers between them. Previous human fMRI studies on resting brain dynamics reported such flattening under artificial conditions such as the administration of psychedelics^7,8^ or transcranial magnetic stimulation.^9^ Further electrophysiological studies in rodents and primates revealed endogenous modulations of cortical potential landscapes during behavioral tasks at higher temporal resolution.^10,11^ However, limitations in the spatiotemporal resolution of activity recordings and/or the complexity of higher vertebrate brains have hindered our understanding of how potential landscape flattening is achieved and controlled at the neuronal circuit level to guide the state transition in a goal-directed manner.

In vertebrates, the cortico-basal ganglia circuit mediates the execution of goal-directed behaviors,^12–17^ and may also mediate state transitions beyond potential barriers by flattening potential landscapes. In this study, by using the adult zebrafish telencephalon as a miniature model of the vertebrate cortico-basal ganglia circuit, we reveal a neural mechanism that temporarily reduces the potential barriers. The teleost telencephalon undergoes a developmental process called eversion, which organizes components similar to those of mammals in mediolaterally inverted positions.^18,19^ In teleosts, the striatum and pallidum are located at the medial and lateral parts of the subpallium, respectively.^20,21^ The central zone of the dorsal pallium (Dc) is homologous to the mammalian isocortex^22,23^ and is involved in higher cognitive functions.^24–27^ We have also shown that these telencephalic regions actually form the conserved cortico-(pallio-)basal ganglia circuit, such as the pallio-striatal pathway, the striatal direct pathway, and the striatal indirect pathway (Fig. 1a, b).^21^ As for the output pathway, in addition to the minor pallidothalamic pathway, there is the fish-specific major shortcut pathway that directly projects from the pallidum (the dorsal entopeduncular nucleus, dEN) to the pallium including the Dc area.^21^ The containment of these main circuit components within about a 1 mm³ cube in the adult zebrafish telencephalon facilitates optical imaging to investigate the circuit’s operating logic.

**Fig. 1:**
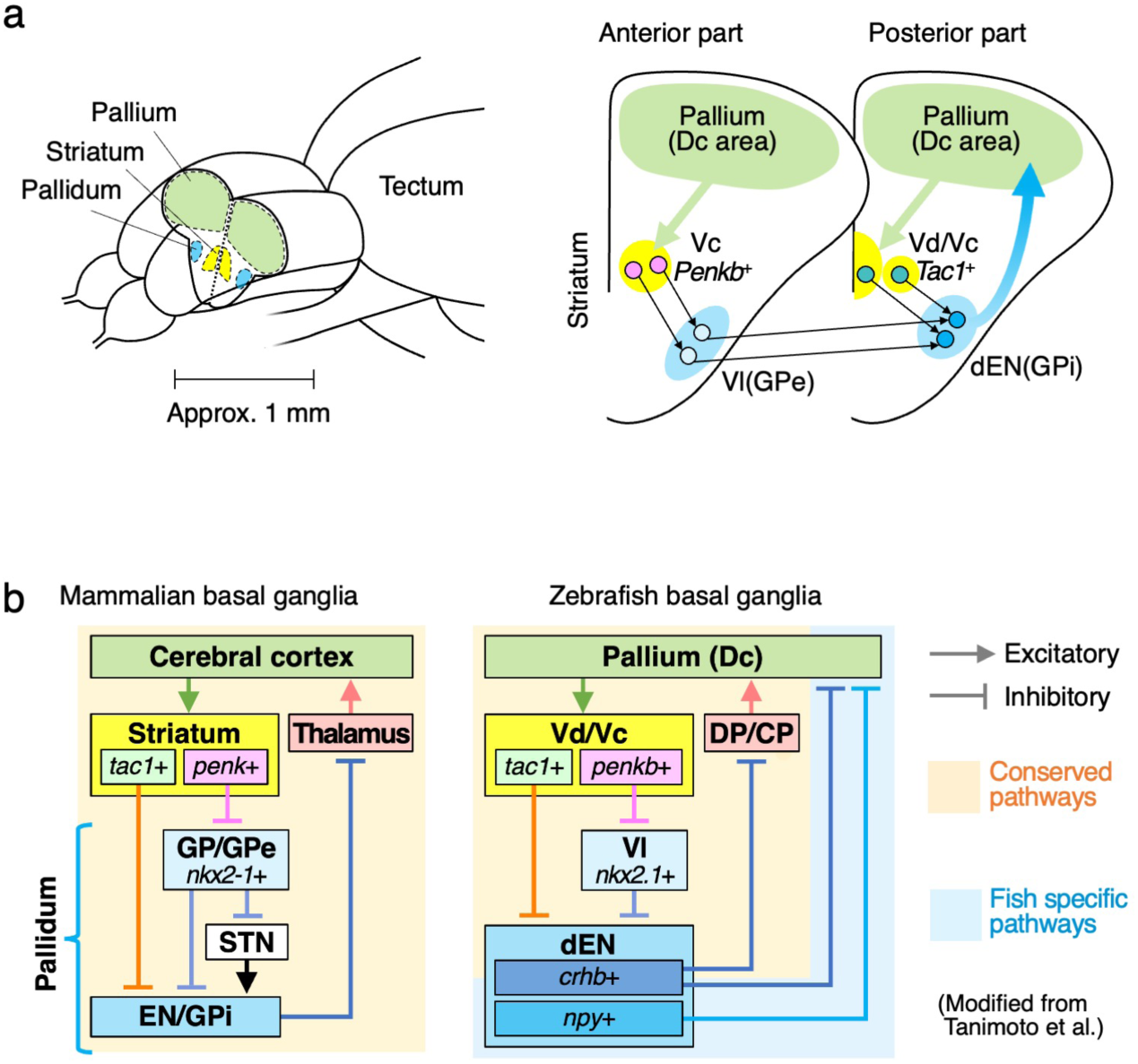
Anatomy of the zebrafish cortico-basal ganglia circuit. **a**, Illustration of the cortico-basal ganglia circuit of zebrafish (modified from Tanimoto et al.^21^). **b,** Schematic diagrams of the mammalian and zebrafish cortico-basal ganglia circuits (modified from Tanimoto et al.^21^). Vd/Vc, dorsal/central nucleus of the ventral telencephalic area; Vl, lateral nucleus of the ventral telencephalic area; dEN, dorsal entopeduncular nucleus; DP/CP, dorsal/central posterior thalamic nucleus. GP, globus pallidus (in rodents); GPe, globus pallidus externus (in primates); STN, subthalamic nucleus; EN, entopeduncular nucleus (in rodents); GPi, globus pallidus internus (in primates).

Moreover, we have previously shown that the Dc acquires various task-related neural ensembles after learning a Go/No-go task in closed-loop virtual reality (VR).^26^ In this task, a head-fixed fish can move forward in a one-dimensional VR space, where the presented scenery moves backward in response to its tail flips (Fig. 2a). After adaptation, the fish learns to escape from the blue region to the red region in Go trials, and to stay in the red region in No-go trials to avoid a mild electric stimulus, which would be given if the fish ends a trial inside the blue region (Fig. 2b). After learning this task, most learner fish generated a “blue/danger”-coding neural ensemble, which encodes danger associated with the blue cue, in their Dc.^26^ One-third of the learner fish additionally generated a “scenery flow prediction error (SFPE)”-coding neural ensemble, which responds to the absence of scenery flow feedback in the open-loop condition (no sensory feedback from the VR).^26^ Although this response is consistent with the idea that brains generate predictions of sensory input,^28^ the SFPE differs from the conventional sensory prediction error, which encodes a mismatch between current and predicted sensory feedback from self-generated motion.^25,29^ The SFPE ensemble, in contrast, also responds to the absence of scenery flow when fish fail to flip their tails during Go trials. This signal can therefore be regarded as another form of prediction error between the current and a desired sensory state.^30,31^ While the fish with a blue/danger-coding Dc ensemble alone escaped from the blue region by moving intermittently, the fish with both blue/danger- and SFPE-coding Dc ensembles showed more efficient escape behavior (Fig. 2c, d).^26^ This improvement in behavioral efficiency suggests that these fish monitor the scenery flow and behave so as to avoid inducing the SFPE.

**Fig. 2:**
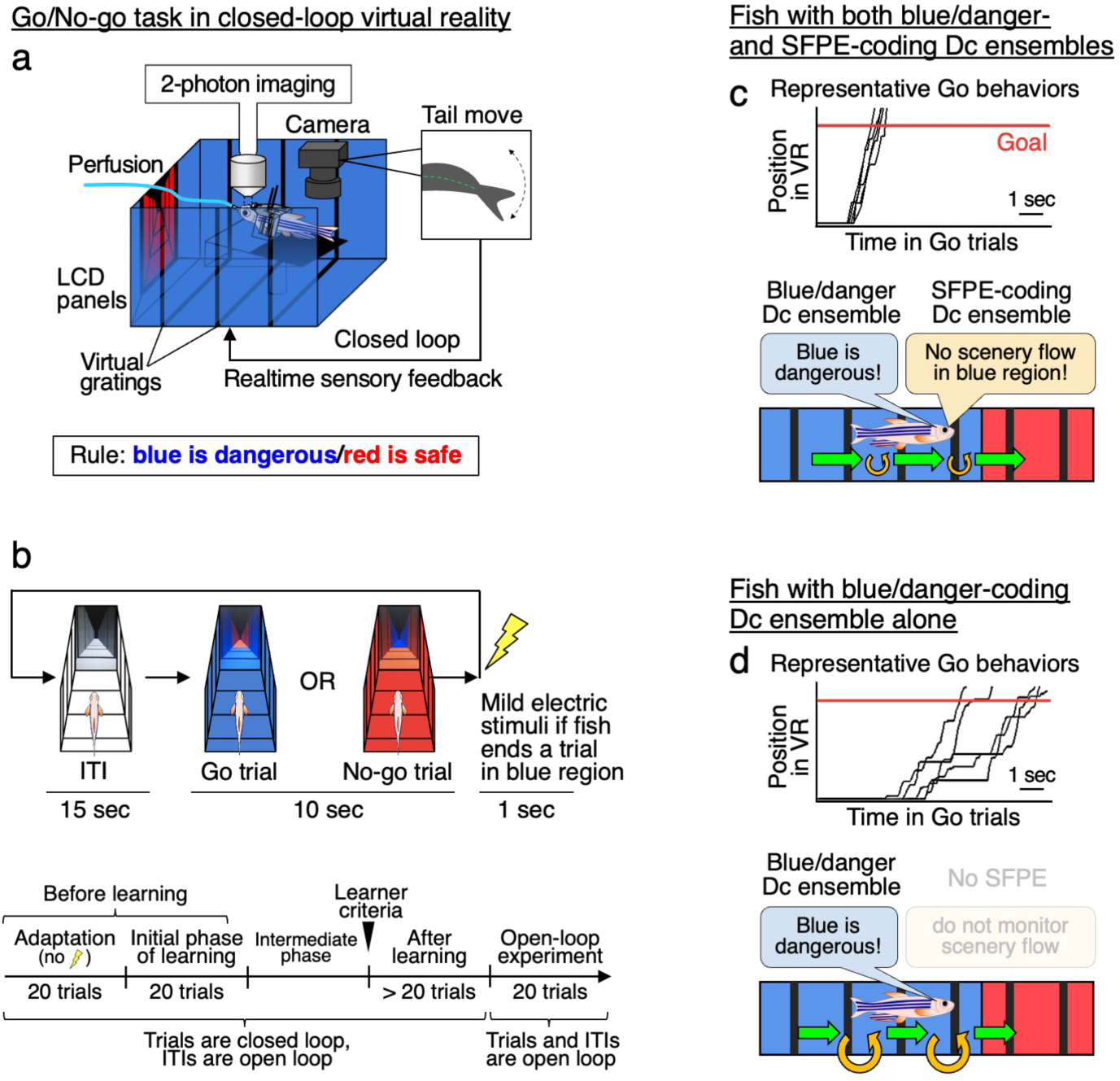
The Go/No-go task in closed-loop virtual reality produced two types of learner fish. **a**, Illustration of the Go/No-go task for adult zebrafish in the closed-loop VR environment (modified from Torigoe et al.^26^). Visual stimuli of a virtual arena were presented on the displays. Tail movements were analyzed in real-time to produce sensory feedback (backward scenery flow) from the VR. **b,** The task design and schedule. After inter-trial intervals (ITIs), a Go or No-go trial was randomly initiated, with a sudden change in background color from white. In Go trials, fish learned to escape from the blue region to the red region within 10 sec. In No-go trials, fish stayed in the red region for 10 sec. Otherwise, fish received a mild electric stimulus as a punishment. **c,** (Top) Representative positional changes during five Go trials and illustration of the behavioral pattern of learner fish with both blue/danger- and scenery flow prediction error (SFPE)-coding Dc ensembles. Position in VR indicates the distance traveled in the virtual arena. (Bottom) Illustration of the efficient escape behavior in this type of fish. Green and orange arrows indicate periods of each move and halt, respectively. **d,** Same as in **c**, but for learner fish with a blue/danger-coding Dc ensemble alone.

Here, we found that the cortico-basal ganglia circuit of zebrafish uses these Dc signals to guide the learned escape behavior by lowering the potential barriers between activity states underlying different behavioral states. After learning the Go/No-go task, their striatal indirect pathway neurons are activated by negative values of the blue-cue and SFPE signals. The dEN output pathway integrates these two signals, and its activation destabilizes ongoing behavioral states. Mathematical modeling revealed that this dEN output can dynamically flatten energy-potential barriers between Dc activity states and facilitate state transitions toward the state with the minimum dEN output activity. Our model explains how pallidal output activity guides state transitions toward choosing an appropriate behavioral state among multiple alternative options.

## Results

### Halt states are destabilized in the fish with the SFPE-coding Dc ensemble

To characterize the different behavioral efficiency of the two types of learner fish (Fig. 2c, d), we divided their escape behavior in the VR during Go trials into forward movement (move state) and halts between move states (halt state). The goal-reaching time was significantly reduced in the fish with both blue/danger- and SFPE-coding Dc ensembles, in comparison with the fish with a blue/danger-coding Dc ensemble alone (Extended Data Fig. 1a, b; original data from Torigoe et al.^26^). This reduction was mostly explained by the different distributions of average halt duration between the two groups (Fig. 3a, b),^26^ and by the shorter halt duration than move duration in the fish with blue/danger- and SFPE-coding Dc ensembles (Fig. 3c, d, top).

**Fig. 3:**
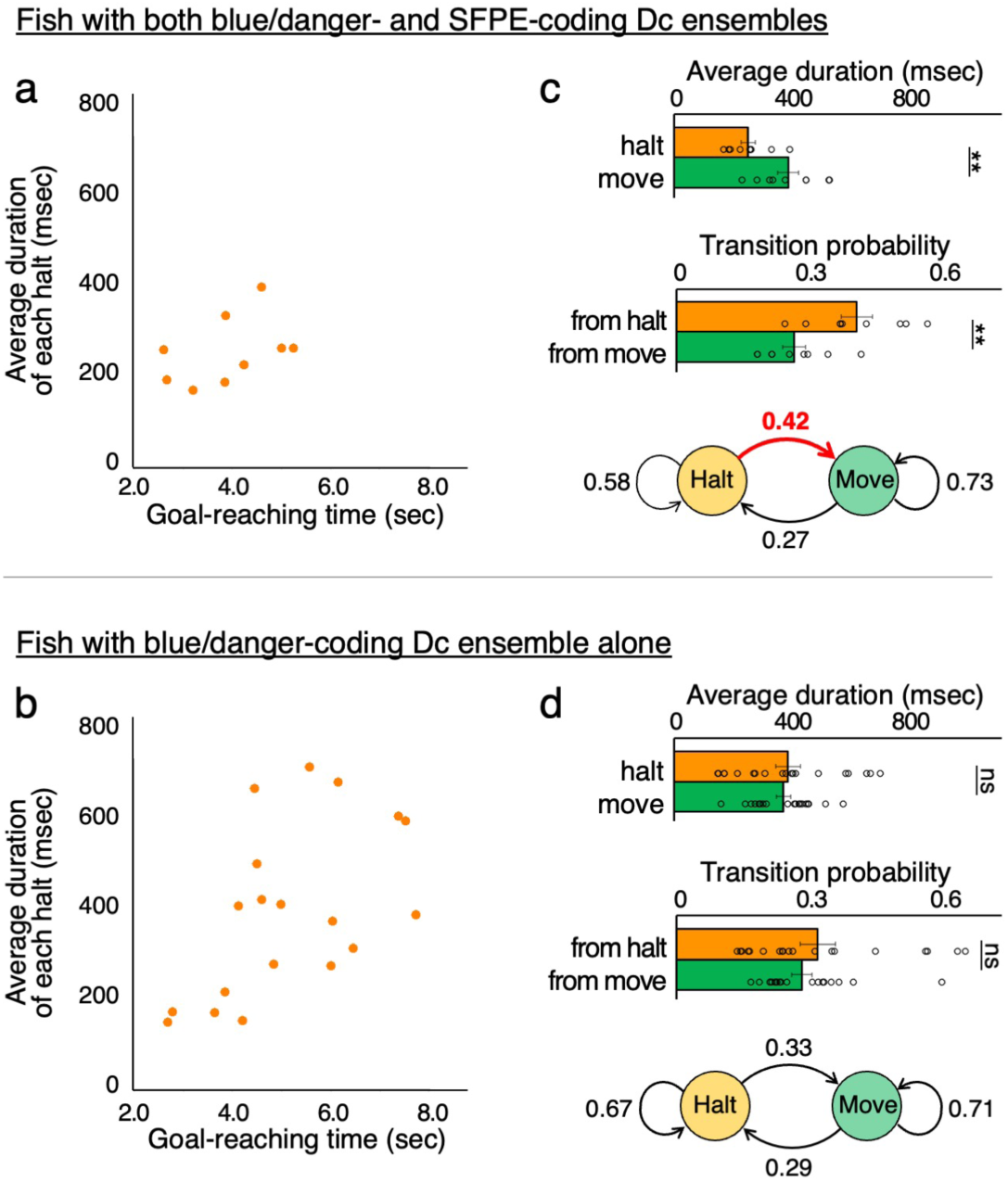
The two types of learner fish have different behavioral transition probabilities. **a**, Distribution of average duration of each halt against goal-reaching time in fish with both blue/danger- and SFPE-coding Dc ensembles. **b,** Same as in **a**, but for fish with a blue/danger-coding Dc ensemble alone. **c,** (Top) Average duration of each halt and move in the fish with both blue/danger- and SFPE-coding Dc ensembles (mean ± s.e.m.; n = 9 fish). Halt duration was shorter than move duration in this type of fish; \*\**P =* 0.0055, two-tailed unpaired *t*-test. (Middle) Transition probabilities during halts and moves for each 100 msec time frame in the same dataset. The transition probability during halts was higher than that during moves; \*\**P =* 0.0055, two-tailed unpaired *t*-test. (Bottom) A state transition model between halts and moves according to the middle panel. The transition probabilities between states are the mean values from the middle panel, and the probabilities of remaining in the same state are obtained by subtracting the transition probability from 1. **d**, Same as in **c**, but for the fish with a blue/danger-coding Dc ensemble alone (n = 19 fish). Similar levels of the halt and move durations and the transition probabilities were observed in this type of fish; ns, not significant, *P =* 0.76 and *P =* 0.44 from top to bottom, two-tailed unpaired *t*-test.

Because the distributions of halt and move durations are approximately linear on a semi-logarithmic plot (*i.e.*, exponential), except for rare, extremely long halts (Extended Data Fig. 1c, d), we approximated the transitions between halts and moves as a Markov process. In this view, short halt duration indicates a high probability of transition out of halts. Accordingly, the fish with blue/danger- and SFPE-coding Dc ensembles showed higher transition probability from the halts than that from the moves, while the fish with a blue/danger-coding Dc ensemble alone did not show such directional inequality in transition probability (Fig. 3c, d, middle). Thus, the behavioral difference between the two learner types is primarily characterized by the different transition probability from the halt state, *i.e*., by how unstable the halt states are (Fig. 3c, d, bottom). This result suggests that halt states are more destabilized and show a greater tendency to transition to move states in the presence of the SFPE-coding Dc ensemble.

### Pallidal output pathway shows uniform activity patterns

To understand the roles of the basal ganglia in this behavior, we performed 2-photon Ca^2+^ imaging of the output pathway from the dEN, the fish homolog of the globus pallidus internus (GPi), during learning of the Go/No-go task. The dEN consists of two GABAergic populations: 84% of the neurons are *npy*^+^ and send direct inhibitory projections to the pallium as the shortcut output pathway, while the minor *crhb*^+^ neurons indirectly provide inhibitory output to the pallium via the thalamus.^21^ Here, we imaged axonal activity of the *npy*^+^ dEN neurons in three different focal planes in the pallium through the skull.^26^ For this, we generated a *Tg(npy:zGFF)* driver line and crossed it with the *Tg(UAS-hsp:zGCaMP6s)* reporter line^32^ to obtain double-transgenic fish *Tg(npy:zGFF);Tg(UAS-hsp:zGCaMP6s)*, which express GCaMP6s in the *npy*^+^ dEN neurons and their axons (Extended Data Fig. 2a). These *npy*^+^ neurons send axonal projections to the broad region of the pallium (Fig. 4a, b).

**Fig. 4:**
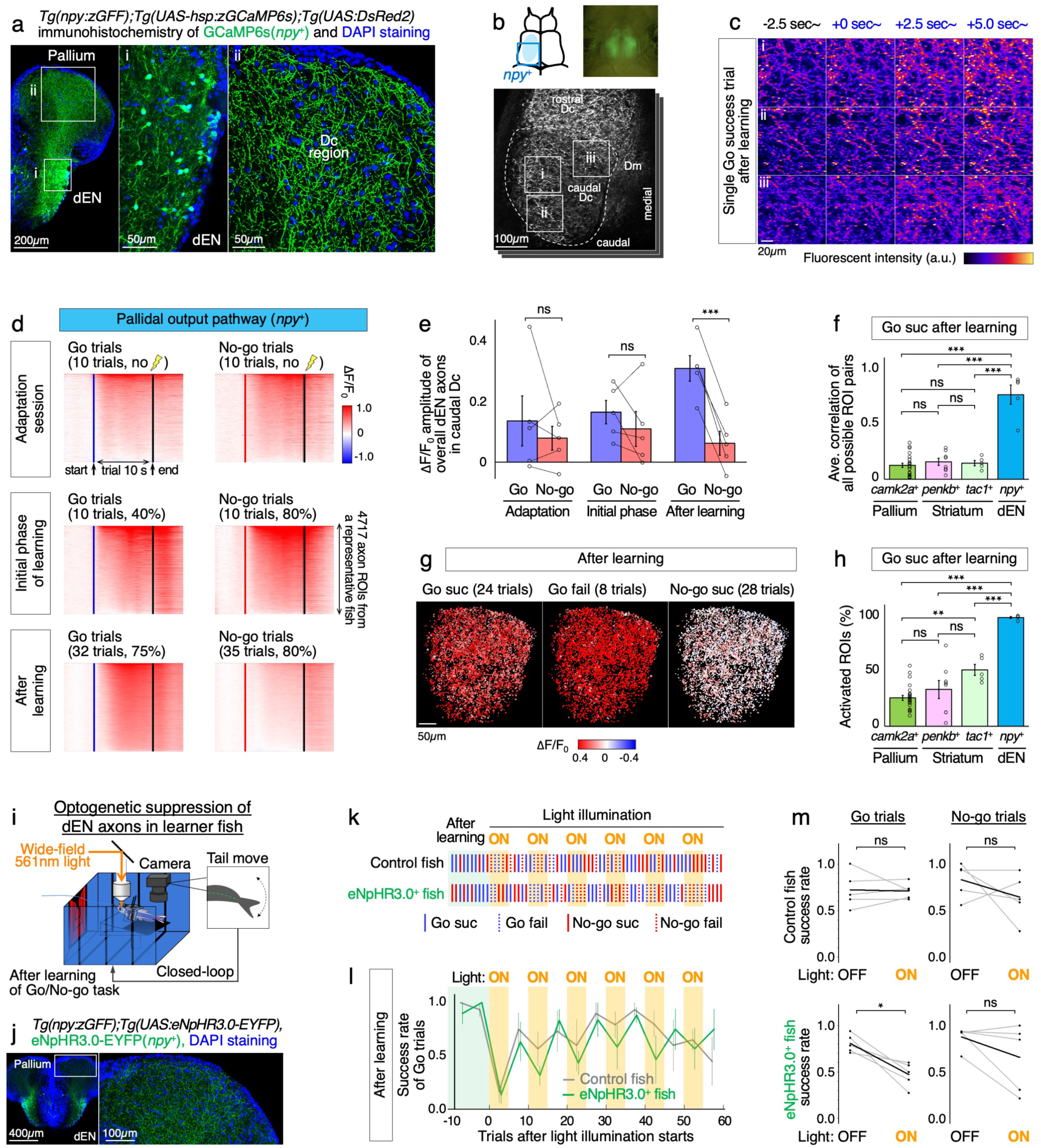
The *npy*^+^ dEN axons broadcast the negative value signal to the Dc for learned escape behavior. **a**, Immunohistochemistry for GCaMP6s (green) expressed in *npy*^+^ dEN neurons and their output axons. A coronal section near the anterior commissure is shown. Insets show the positions of the panels i and ii. **b,** (Top left) Illustration of the approximate position of the imaging view-field, focusing on *npy*^+^ dEN axons in the Dc (cyan inset). (Top right) A fluorescent dissection microscopy image of a fish used for this imaging. (Bottom) Two-photon imaging of *npy*^+^ dEN output axons in the pallial Dc region through the skull. Dashed line indicates the caudal Dc. Insets show the positions of **c**. Dm, medial zone of the dorsal pallium. **c,** Uniform activation of dEN axons in the three different areas of the caudal Dc during one representative Go success trial after learning. Time-averaged fluorescent images for each 2.5 sec bin starting from the indicated time points are shown. Presentation of the blue cue starts at 0 sec. **d,** Learning-dependent activity changes of *npy*^+^ dEN axons in the Dc area during Go and No-go trials. Trial-averaged ΔF/F_0_ patterns (heatmaps) from one representative fish are shown. The number of trials used for trial-averaging is indicated above each heatmap, along with the Go and No-go success rates during adaptation, the initial phase of learning, and the after-learning phase. The axonal ROIs are sorted according to their time-averaged response amplitudes during trials in each heatmap. Blue and red vertical lines indicate the onset of blue and red cues, respectively. Black vertical lines indicate the end of the trial. **e,** Activity from overall dEN axons in the Dc of multiple fish (mean ± s.e.m.; n = 5 fish). The ΔF/F_0_ amplitudes in Go trials became higher than those in No-go trials after learning; \*\*\**P* < 0.0001, two-tailed paired *t*-test. **f,** Averaged temporal correlation coefficients between all possible ROI pairs within each fish for *camk2a*^+^ Dc excitatory neurons (n = 24 fish), *penkb*^+^ striatal neurons (n = 8 fish), *tac1*^+^ striatal neurons (n = 5 fish), and *npy*^+^ dEN axons (n = 5 fish). Go success trials after learning were used. \*\*\**P <* 0.0001, one-way ANOVA with Tukey’s HSD post hoc test. **g,** Spatial distribution of ΔF/F_0_ amplitudes across dEN axonal ROIs during Go success, Go failure, and No-go success trials after learning. Distributions of three focal planes from the same representative fish as **d** were overlaid by maximum intensity projection. **h,** Proportion of activated ROIs (ΔF/F_0_ > 5%) during Go success trials after learning. The same dataset as **f**. Almost all the dEN axonal ROIs were activated; \*\*\**P <* 0.0001, \*\**P =* 0.0033, one-way ANOVA with Tukey’s HSD post hoc test. **i,** Illustration of the optogenetic suppression experiment of dEN axons. **j,** Immunohistochemistry for eNpHR3.0-EYFP (green) expressed in *npy*^+^ dEN output neurons. Inset shows the positions of the right panel. **k,** Behavioral performance of representative control and eNpHR3.0^+^ fish during repeated light ON (orange shading) and OFF (no shading) blocks for 5 trials. Before the light illumination, the fish were already in a learned state (green shading). Dotted bars indicate failure trials and solid bars indicate successful trials. **l,** Temporal changes in the success rate of Go trials during each light ON and OFF block for control (mean ± s.e.m.; n = 5 fish) and eNpHR3.0^+^ (n = 5 fish) conditions in 5-trial bins. The horizontal axis shows the number of trials completed after the onset of the first light illumination. **m,** Statistics of the success rate of Go and No-go trials during all the light ON and OFF trials except for the first ON block. Gray lines indicate each fish (the same dataset as **l**), and a thick black line indicates the mean. The success rate of Go trials was significantly reduced in eNpHR3.0^+^ fish during light ON; \**P =* 0.017, two-tailed paired *t*-test.

These *npy*^+^ dEN axons showed uniform activation in the caudal Dc area after learning (Fig. 4c, Supplementary Videos 1 and 2). Trial-averaging of each axon’s activity pattern revealed that most of the axons in this area showed higher activation in Go trials than in No-go trials after learning (Fig. 4d, e). Detailed comparison across trial types (Extended Data Fig. 2b, c) showed that these axons were active during occasional tail movement in Go success trials in the initial phase of learning (see Discussion for interpretation of this result). In addition, in Go failure trials, response amplitudes were significantly increased after learning compared with the initial phase of learning (Extended Data Fig. 2c). This suggests that, after learning, these axons responded not only during tail movement but also to the blue cue itself even without proper tail movement. No-go failure trials were rare and therefore not analyzed, possibly reflecting the inherent tendency of head-fixed fish to stay.^26^ Activity during No-go success trials was not significantly affected by learning (Extended Data Fig. 2c). During Go success trials after learning, the activity patterns were highly correlated among axon branches (Fig. 4f), and almost all of them were co-activated in a spatially non-biased manner (Fig. 4g, h), in contrast to the activation patterns of the Dc excitatory neurons (as we previously reported in Torigoe et al.^26^) or the striatal neurons (as reported here; see below). This activation pattern adaptively changed by rule switching from the original rule [blue is dangerous/red is safe] to the reversed rule [red is dangerous/blue is safe], *i.e.,* these axons became preferentially activated to the dangerous red cue after reversal learning (Extended Data Fig. 3a-d). These results indicate that the dEN output pathway adaptively responds to a color cue paired with the punishment and uniformly broadcasts the information to the caudal Dc area of the pallium.

### Pallidal output suppression impairs learned escape behavior

To test whether this learning-dependent dEN activity affects the learned escape behavior, we optogenetically suppressed the dEN axons after learning the Go/No-go task in the original rule using the enhanced halorhodopsin eNpHR3.0.^33^ We generated *Tg(npy:zGFF);Tg(UAS:eNpHR3.0-EYFP)* fish and suppressed the output activity by wide-field illumination of 561 nm light onto the bilateral pallium (Fig. 4i, j). The light stimulation was turned ON and OFF in alternating blocks of 5 trials after learning (Fig. 4k). In the first light ON trials, both control and eNpHR3.0^+^ fish showed a reduction in the success rate of Go trials, likely due to a startle response to the unaccustomed light leaking into their eyes (Fig. 4l). However, in the subsequent trials, only the eNpHR3.0^+^ fish showed a decreased Go success rate in the light ON trials in comparison with the light OFF trials (from 0.81 to 0.47, *P* = 0.017, Fig. 4m). Since this reduction of the Go success rate was reversible and was restored quickly after turning off the light, the output from the dEN to the pallium plays a significant role in the proper execution of the learned escape behavior.

### Pallidal output activation destabilizes ongoing behavioral states

We next investigated the function of the dEN output activity. These dEN neurons are GABAergic.^21^ The established view of the basal ganglia output function is to inhibit actions.^34^ However, here the dEN output pathway is uniformly activated for the learned escape behavior, implying that it rather facilitates action execution. To reveal how this uniform activation affects behavior, we optogenetically activated the dEN output pathway using a transgenic line *Tg(UAS:ChrimsonR-tdTomato)*^35^ for expression of the red-shifted channelrhodopsin ChrimsonR^36^ in the dEN axons (Fig. 5a). In naïve *Tg(npy:zGFF);Tg(UAS:ChrimsonR-tdTomato);Tg(UAS-hsp:zGCaMP6s)* fish (Fig. 5b), uniform light illumination of dEN axons in the unilateral pallium activated the dEN axons, particularly in the caudal Dc area (Fig. 5c, Supplementary Video 3). During this activation, tail movement was recorded and classified into move states and stay states by the same detection threshold used for the VR (6 degrees, Fig. 5d). In this analysis, we included intermittent short periods (< 1 sec) with no tail flip as parts of one episode of move state when they occur between tail-flipping episodes. In the control fish only carrying *Tg(npy:zGFF);Tg(UAS-hsp:zGCaMP6s)*, neither stay-to-move nor move-to-stay transition was affected by light illumination under the head-fixed condition (Fig. 5d, e, top panels). In the ChrimsonR^+^ fish, which additionally carry *Tg(UAS:ChrimsonR-tdTomato)*, the behavioral transition events between the move and stay states significantly increased in response to light illumination, both in stay-to-move and move-to-stay directions (Fig. 5d, e). These results indicate that uniform activation of the dEN output destabilizes ongoing behavioral states and facilitates state transitions.

**Fig. 5:**
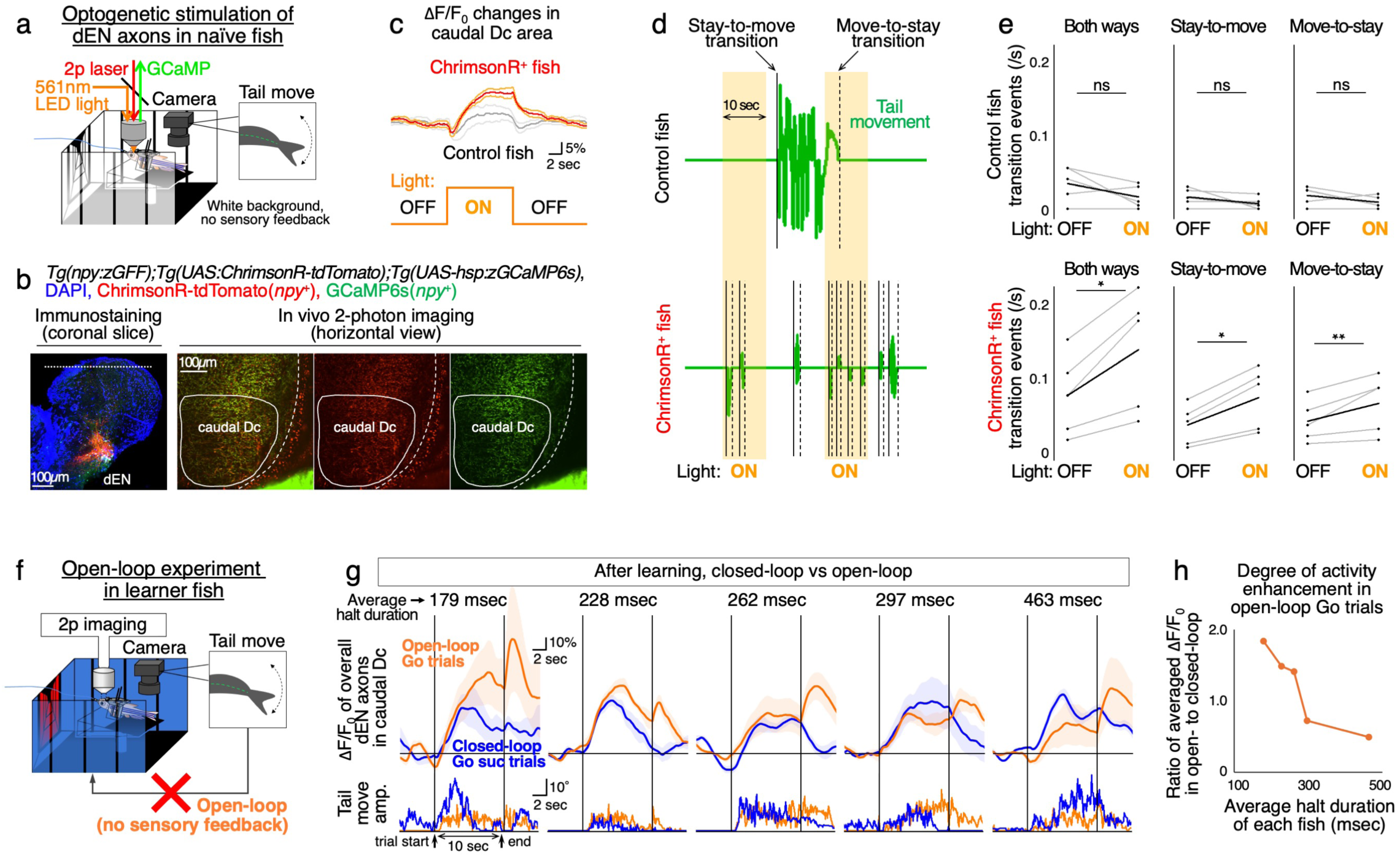
The *npy*^+^ dEN axons destabilize ongoing behavioral states and integrate blue-cue and SFPE signals. **a**, Illustration of the optogenetic stimulation experiment of the dEN axons in naïve fish. Tail movement and simultaneous neural activity of the stimulated dEN axons were recorded during 20 light stimulations in a constant white background color. **b,** Expression of ChrimsonR-tdTomato (red) and GCaMP6s (green) in *npy*^+^ dEN output neurons. (Left) Immunohistochemistry for ChrimsonR-tdTomato and GCaMP6s (coronal slice). Dotted line indicates the approximate dorso-ventral level where the horizontal sections in the right panels were made. (Right) 2-photon imaging of the ChrimsonR-tdTomato and GCaMP6s in vivo (horizontal view). Solid lines indicate the caudal Dc. Dashed lines indicate the outline of the brain. **c,** ΔF/F_0_ changes in the caudal Dc area during light stimulation for control (n = 5 fish, gray line) and ChrimsonR^+^ fish (n = 5 fish, red line). The line graph shows mean traces across fish, with lighter lines indicating mean ± s.e.m. traces. Before averaging across fish, all 20 light stimulations were trial-averaged. **d,** Representative tail movement patterns of control and ChrimsonR^+^ fish. Behavioral transitions were detected as either stay-to-move (solid line) or move-to-stay (dashed line) transitions. **e,** Statistics of transition events (/sec) in both ways, stay-to-move, and move-to-stay transitions during light ON and OFF phases. Gray lines indicate each fish (the same dataset as **c**), and a thick black line indicates the mean. The transitions were increased by light illumination; \**P =* 0.014, \**P =* 0.039, \*\**P =* 0.0062 from left to right, two-tailed paired *t*-test. **f,** Illustration of the open-loop experiment after learning the Go/No-go task. **g,** Activity patterns of overall dEN axons in the Dc during open-loop Go trials. Trial-averaged ΔF/F_0_ patterns (mean ± s.e.m.) across 5 closed-loop Go success trials (blue lines) and 5 open-loop Go trials (orange lines), just before and after initiation of the open-loop experiment. Data from the same 5 fish as the npy^+^ dEN data in Fig. 4e-h are shown, from the one with the shortest halt duration (leftmost panel) to the one with the longest halt duration (rightmost panel). The simultaneous tail movement amplitudes are shown below in the same line colors as the top panels. Horizontal lines indicate ΔF/F_0_ = 0. Scale bars, 10% and 2 sec for ΔF/F_0_; 10° and 2 sec for tail movement. **h,** Ratio of time-averaged activity during the open-loop Go trials to that during the closed-loop Go success trials. This ratio is plotted against the average halt duration for each fish (n = 5 fish).

### Pallidal output pathway integrates blue/danger and SFPE signals

We further found that the dEN integrates multiple signal components. In our closed-loop VR system, backward scenery flow is generated when fish flip their tails during trials. Here, we also performed an open-loop experiment after learning (Fig. 5f), where the backward scenery flow is not delivered from the VR by the tail movement and therefore the SFPE signal is strongly induced.^26^ The dEN axons showed enhanced activity during open-loop Go trials compared with activity during closed-loop Go success trials, particularly in the fish with shorter halt duration (Fig. 5g, left panels). In contrast, in the fish with longer halt duration, the dEN axons were activated just by the blue-cue presentation at similar levels during the open- and closed-loop Go trials (Fig. 5g, right panels). We quantified the degree of this enhancement from the axonal activity traces and found that it gradually decreased as the average halt duration of the fish increased (Fig. 5h). These results suggest that the enhanced component of dEN activity on top of the blue-cue component in the open-loop condition reflects the SFPE signal.

Our observation on the Dc showed that fish with blue/danger- and SFPE-coding Dc ensembles have shorter halts (see Fig. 3c, d).^26^ Considering that the dEN receives information derived from the Dc through the pallio-striatal-dEN pathway (see Fig. 1), the enhancement of the dEN output activity may be caused by integration of blue/danger- and SFPE-signals from the Dc in fish having shorter halts. In contrast, in fish having longer halts, the dEN activity may be affected by the blue/danger signal alone, as the SFPE signal is lacking in the Dc of such fish. Taken together, these observations suggest that the dEN neurons integrate the blue/danger and SFPE signals from the Dc.

### The striatal indirect pathway mediates the negative value signals

To elucidate how afferent neural circuits control the dEN activity, we next imaged striatal indirect- and direct-pathway neurons, which are *penkb*^+^ and *tac1*^+^, and are located in the anterior and posterior parts of the fish striatum, respectively (see Fig. 1a, b).^21^ We previously showed that both of these striatal populations receive projections from the Dc, and provide anatomical pathways from the Dc to the dEN.^21^ The *penkb*^+^ indirect-pathway neurons project to the lateral nucleus of the ventral telencephalic area (Vl), fish globus pallidus externus (GPe), which in turn projects to the dEN, forming a net excitatory pathway to the dEN due to the two inhibitory connections (Extended Data Fig. 4a-c).^21^ The *tac1*^+^ direct-pathway neurons directly project to the dEN.^21^

To image striatal indirect- and direct-pathway neurons, we generated *Tg(penkb:zGFF);Tg(UAS-hsp:zGCaMP6s)*and *TgBAC(tac1:GAL4VP16);Tg(UAS-hsp:zGCaMP6s)* fish, respectively (Fig. 6a, b).^21,32^ For 2-photon imaging of the striatum, which is located approximately 400 µm below the surface of the telencephalon, we used a cranial window covered with a microsheet of Teflon (Fig. 6c, Extended Data Fig. 5a-b, Supplementary Videos 4-7), which is highly transparent due to the refractive index similar to that of water.^37,38^ Before learning, some indirect- and direct-pathway neurons were active during Go success trials (Fig. 6d-f, Extended Data Fig. 6). After learning, we observed a significant increase in the number of indirect-pathway neurons activated during Go failure trials (Fig. 6d, e). This suggests that a part of the indirect-pathway neurons acquired responsiveness to the blue cue by learning. Direct-pathway neurons also showed a similar tendency to the indirect pathway (Fig. 6f, Extended Data Fig. 6).

**Fig. 6:**
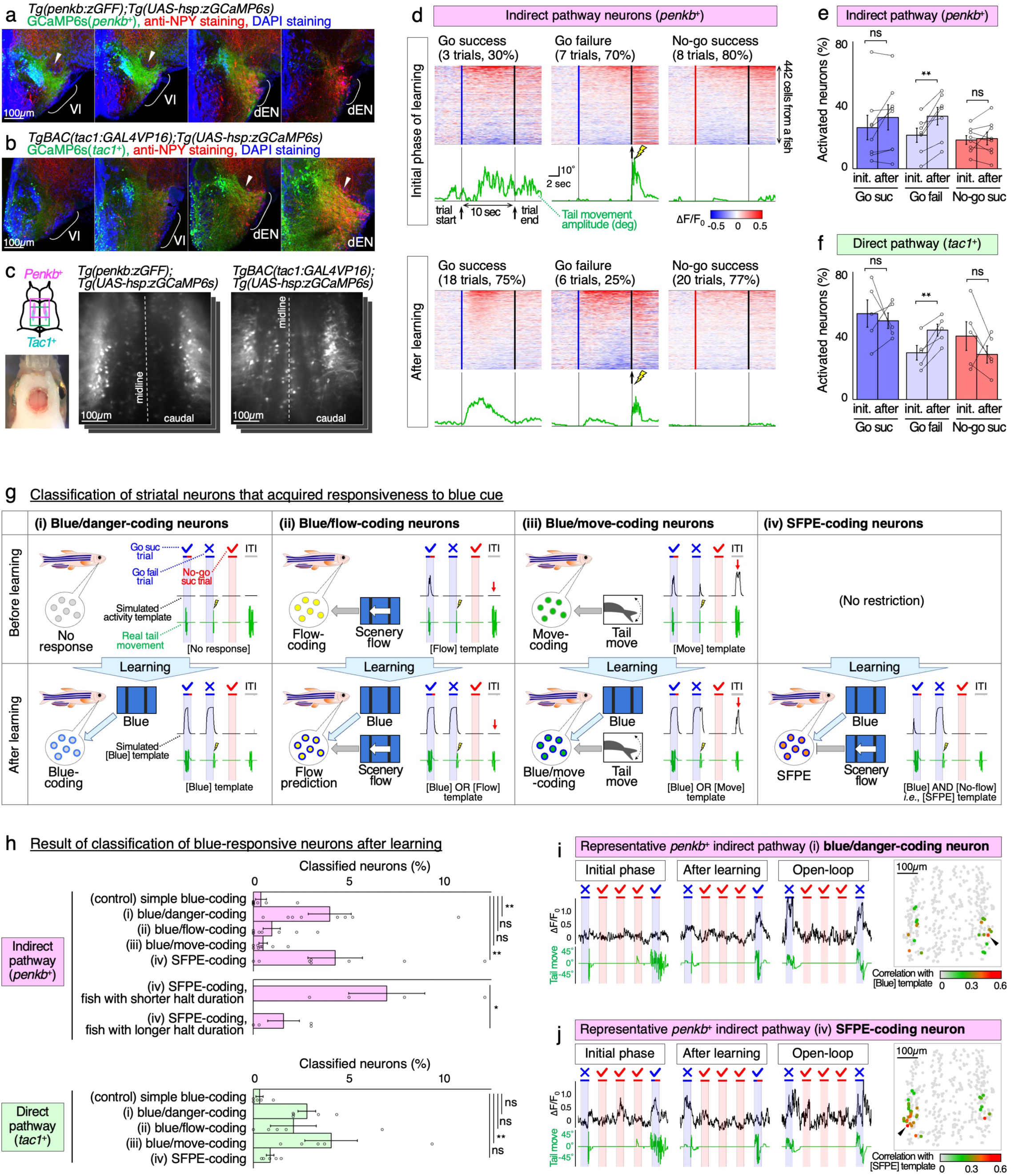
The striatal indirect pathway neurons encode the blue-cue and SFPE signals after learning. **a**, Immunohistochemistry for GCaMP6s (green) expressed in *penkb*^+^ indirect-pathway striatal neurons. Four successive coronal sections from the anterior (left panels) to the posterior (right panels) parts of the telencephalon are shown. The anterior two sections contain the NPY⁻ Vl region, and the posterior two sections contain the NPY^+^ dEN region (red). Brackets indicate either the Vl or dEN region. Arrowheads indicate a projection to the Vl region. **b,** Same as in **a**, but for *tac1*^+^ direct-pathway striatal neurons. Arrowheads indicate a projection to the dEN region. **c,** (Left top) Illustration of the approximate positions of the imaging view-fields, focusing on *penkb*^+^ neurons (magenta inset) and *tac1*^+^ neurons (light green inset). (Left bottom) A dissection microscopy image of a fish with a cranial window. (Right) Two-photon imaging of striatal neurons using the cranial windows. **d,** Learning-dependent activity changes of *penkb*^+^ striatal neurons during Go success, Go failure, and No-go success trials. Trial-averaged ΔF/F_0_ patterns (heatmaps) and simultaneous tail movement amplitudes (green lines) from one representative fish are shown. The number of trials used for trial averaging is indicated above each heatmap, along with the Go success, Go failure, and No-go success rates during the learning phase. The neurons are sorted according to their time-averaged response amplitudes during the trials in each heatmap. Lightning marks indicate the timing of mild electric stimuli. Other symbols, labels, and annotations are the same as in Fig. 4d. **e,** Learning-dependent changes in proportions of activated *penkb*^+^ neurons (ΔF/F_0_ > 5%) from multiple fish (mean ± s.e.m.; n = 8 fish). The activated proportion in Go failure trials increased after learning; \*\**P =* 0.0028, two-tailed paired *t*-test. **f,** Same as in **e**, but for *tac1*^+^ direct-pathway striatal neurons (n = 5 fish). The activated proportion in Go failure trials increased after learning; \*\**P =* 0.0079, two-tailed paired *t*-test. **g,** Four hypothetical response types of striatal neurons that acquire blue responsiveness through learning. The four response types (i-iv) are arranged from left to right, with the top and bottom rows showing before and after learning, respectively. In each panel, the left illustration shows the events to which the neuron type responds (blue cue, scenery flow, or tail movement), and the right graph shows a representative simulated activity template (black line; see Methods) based on the real tail movement of the fish (green line). (i) Blue/danger-coding neurons are initially non-responsive and become blue-responsive. (ii) Blue/flow-coding neurons are initially scenery flow-responsive and become additionally blue-responsive. Since scenery flow is not delivered during ITIs, these neurons do not respond to tail movement during ITIs (red arrow). (iii) Blue/move-coding neurons are initially move-responsive and become additionally blue-responsive. These neurons respond to tail movement during ITIs (red arrow). (iv) SFPE-coding neurons become responsive to the absence of scenery flow in the blue region. Blue and red shading indicate periods during Go and No-go trials, respectively. The blue or red color bars at the top indicate color cues presented. Lightning marks indicate the timing of mild electric stimuli. **h,** (Top) Proportion of classified *penkb*^+^ neurons into the five response types (mean ± s.e.m. across n = 8 fish). Significantly more (i) blue/danger-coding neurons and (iv) SFPE-coding neurons were detected than (control) simple blue-coding neurons; \*\**P =* 0.0088, \*\**P =* 0.0049 from top to bottom, Dunnett’s test with the alternative hypothesis being greater than the control. (Middle) Comparison of the proportion of classified *penkb*^+^ SFPE-coding neurons between fish with shorter halt duration (below the median halt duration, n = 4 fish) and fish with longer halt duration (above the median halt duration, n = 4 fish). More SFPE-coding neurons were detected in the fish with shorter halt duration; \**P =* 0.046, two-tailed unpaired *t*-test. (Bottom) Same as in (top), but for *tac1*^+^ neurons (n = 5 fish). Significantly more (iii) blue/move-coding neurons were detected than the control neurons; \*\**P =* 0.0096, Dunnett’s test with the alternative hypothesis being greater than the control. **i,** Activity of a representative *penkb*^+^ (i) blue/danger-coding neuron (initially non-responsive and became blue-responsive after learning). For each phase, 5 trials are shown, with the same symbols and annotations as in **g**. The rightmost panel indicates a distribution of *penkb*^+^ (i) neurons in the representative fish. The color code indicates the correlation coefficient with the corresponding template after learning. Arrowhead indicates the position of the representative neuron. Gray dots indicate all the neurons other than (i) neurons. **j,** Same as in **i**, but for a representative *penkb*^+^ (iv) SFPE-coding neuron. As defined in the template, the SFPE neuron responded to the absence of backward scenery flow in the blue region after learning, due to the fish’s failure to flip its tail, or due to the open-loop experiment (no scenery flow feedback from the VR by the tail movement). Other symbols, labels, and annotations are the same as in **i**.

Next, we classified the striatal neurons that acquired responsiveness to the blue cue (Fig. 6g). We generated the following hypotheses based on each neuron’s response property before learning, specifically to the blue cue, scenery flow, tail movement, or none of these. (i) Blue/danger-coding neurons, which are initially non-responsive and become blue-responsive after learning; (ii) Blue/flow-coding neurons, which are initially scenery flow-responsive and become additionally blue-responsive after learning, and may serve to predict scenery flow; (iii) Blue/move-coding neurons, which are initially tail movement-responsive (both during trials and ITIs; in ITIs, tail movement does not produce scenery flow; see Fig. 6g red arrow) and become additionally blue-responsive after learning. In addition, we included (iv) SFPE-coding neurons, which respond to the absence of scenery flow in the blue region regardless of tail movement after learning (see Fig. 2c). As a control condition, simple blue-coding neurons, which are blue-responsive both before and after learning, were also included. Based on these assumed response types, we simulated activity templates from the measured behavioral data for each fish (Fig. 6g; see Methods). Then, each striatal neuron was classified into at most one of the response types according to the criteria defined for its correlation coefficient with the template before and after learning (Extended Data Fig. 7a, b; see Methods). Examples of classified neurons with different correlation coefficients after learning are shown in Extended Data Fig. 7c-h. In comparison with the control neurons, the indirect pathway contained significantly more (i) blue/danger-coding neurons and (iv) SFPE-coding neurons (Fig. 6h-j). More SFPE-coding neurons were detected from the indirect pathway in the fish with shorter halts than in the fish with longer halts (Fig. 6h middle). These observations recapitulate our previous observations in the Dc of the two types of learner fish (see Fig. 2c, d),^26^ and suggest that the observed activities in the indirect pathway mediate the blue/danger- and SFPE-signals derived from the Dc. Since both of these signals are paired with punishment, the observed activities in the indirect pathway reflect their negative values.

In contrast to the control neurons, the direct pathway contained significantly more (iii) blue/move-coding neurons, which were initially move-responsive and became additionally blue-responsive (Fig. 6h bottom, Extended Data Fig. 7g-h). Considering that tail movement results in the successful outcome in Go trials, this activity in the direct pathway may encode the positive value of the tail movement in the presence of the blue cue. We will report further characterization of the direct pathway elsewhere.

### A neural circuit model of the fish basal ganglia

Our observations have led us to propose a circuit model of the zebrafish basal ganglia after learning (Fig. 7a). In this model, the *penkb*^+^ indirect-pathway neurons become responsive to negative values of the blue/danger- and SFPE-coding signals. These two signals from the indirect pathway neurons are then transmitted to the dEN as net excitatory inputs due to the two inhibitory connections of the indirect pathway via the Vl (see Extended Data Fig. 4). The *npy*^+^ dEN output neurons integrate these inputs as a sum of the two components, and uniformly broadcast the integrated negative value signal to the Dc area through their extensively branching axons. This output activity from the dEN causes destabilization of ongoing behavioral states. The teleost pallium consists mostly of excitatory neurons and partly of inhibitory interneurons, as in the case of mammalian cortex (Extended Data Fig. 8a, b).^21,24,39,40^ Through their extensively branching axons, the *npy*^+^ dEN neurons appear to project to almost all the Dc neurons, including the inhibitory interneurons in the Dc area (Extended Data Fig. 8c-f).

**Fig. 7:**
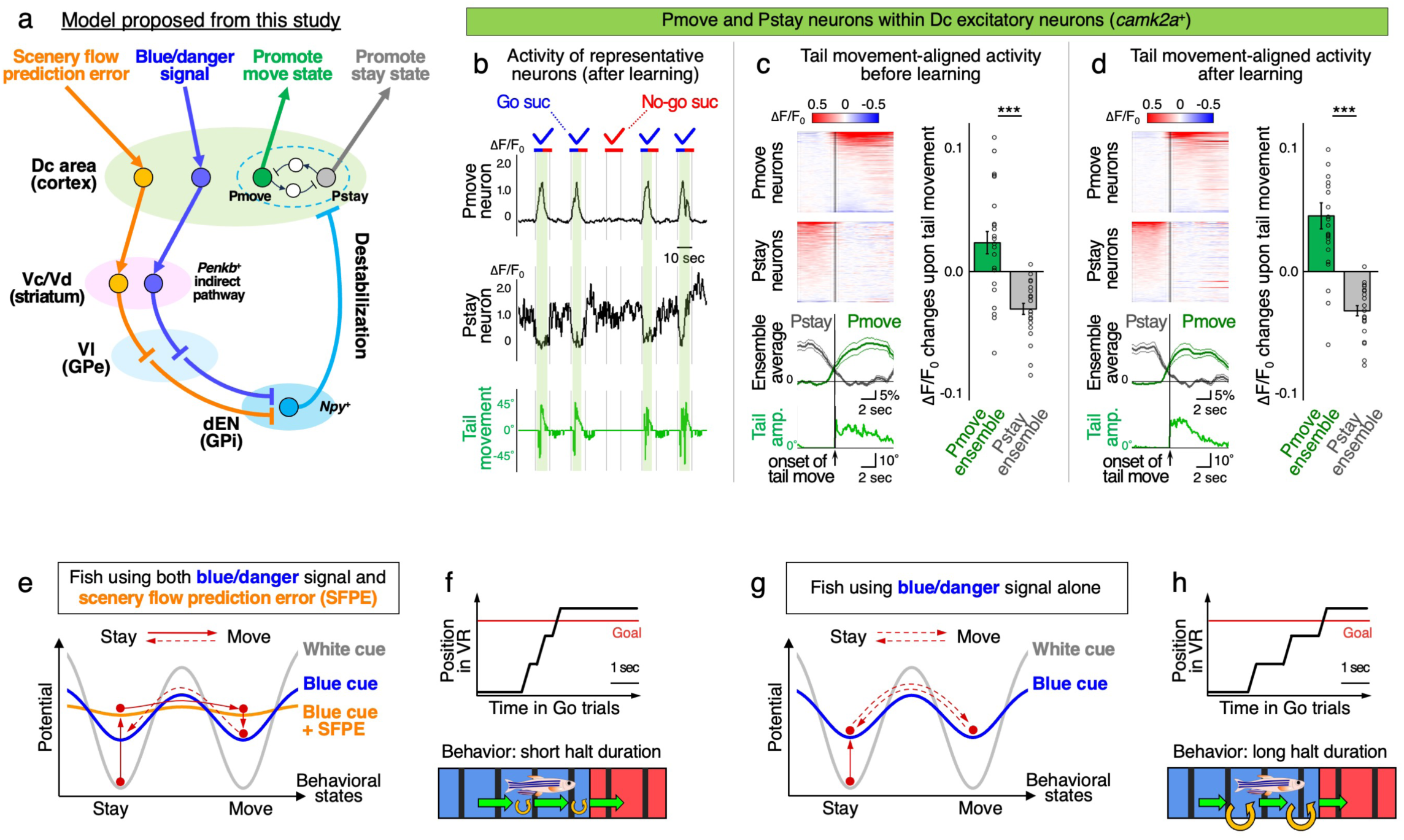
The circuit model of the zebrafish basal ganglia for goal-directed state transition. **a**, Schematic diagram of a putative operational neural circuit, which is established after learning the Go/No-go task. **b,** Representative Dc excitatory neurons putatively promoting move states (Pmove neurons) or promoting stay states (Pstay neurons). Activity patterns of a Pmove neuron and a Pstay neuron (black lines) along with simultaneous tail movement (green line) from one representative fish are shown (original data from Torigoe et al.^26^). Green shading indicates time windows during activation of the Pmove neuron. **c,** Peri-event time histograms of Pmove and Pstay activity before learning. (Left top) Heatmaps of event-averaged ΔF/F_0_ patterns across 19 tail movement events of all the Pmove and Pstay neurons from one representative fish. Vertical lines indicate the timing of tail movement onset. Pmove and Pstay activities were normalized by F_0_ during pre- and post-event 5 sec, and sorted according to their time-averaged response amplitudes during post- and pre-event 5 sec, respectively. (Left bottom) Ensemble-averaged ΔF/F_0_ patterns (mean ± s.e.m.) across all the Pmove neurons (n = 144 neurons, dark green line) and all the Pstay neurons (n = 118 neurons, gray line) from the same representative fish. Light green line indicates simultaneous event-averaged tail movement amplitudes. (Right) ΔF/F_0_ changes of the Pmove and Pstay neurons between pre-event (5 sec) and post-event (5 sec) periods across multiple fish (n = 24 fish); \*\*\**P <* 0.0001, two-tailed paired *t*-test. **d,** Same as in **c**, but for after learning (28 tail movement events). (Left) The same representative fish, and the same sets of Pmove and Pstay neurons as in **c,** left panels. (Right) The same dataset as in **c,** right panel (n = 24 fish); \*\*\**P <* 0.0001, two-tailed paired *t*-test. **e,** Hypothetical potential landscape changes during Go success trials after learning in fish using both blue/danger- and SFPE-signals. Dashed lines indicate less likely transitions compared with the solid lines. **f,** Illustration of efficient escape behavior with shorter halts in this type of fish. **g-h,** Same as in **e-f**, but for fish using blue/danger signal alone, which showed less efficient escape behavior with prolonged halts.

By re-analyzing our previous recording data on the *camk2a*^+^ Dc excitatory neurons,^26^ we found neurons whose activation was accompanied by increased probability of tail movement (Fig. 7b, > 1.5-fold increase), and we hereafter refer to them as Pmove neurons. There were also neurons (hereafter Pstay neurons), whose activation was accompanied by decreased probability of tail movement (Fig. 7b). Peri-event time histograms aligned to the initiation of tail movement showed increased and decreased levels of activity of the Pmove and Pstay neurons at the onset of tail movement, respectively, irrespective of before and after learning (Fig. 7c, d). For such antagonistic activities of two neuronal populations, one of the simplest circuit models is that each population mutually inhibits the other via the local inhibitory interneurons, while forming separate neural ensembles via recurrent excitatory connections (see Fig. 7a). According to the model, which incorporates these observations, activating GABAergic dEN axons is expected to destabilize ongoing behavioral states by suppressing mutual inhibition between the Pmove and Pstay ensembles.

### The operating logic of the fish basal ganglia for goal-directed behavior

We next examined how the identified circuit could produce the different behavioral patterns of the two types of learner fish (see Fig. 2c, d). We hypothesized that the firing response of the dEN output encodes the sum of blue/danger- and SFPE-signals with much faster dynamics than the observed calcium responses, which are limited by the temporal resolution of calcium imaging. Under this hypothesis, the dEN could quickly destabilize ongoing neural and behavioral states as follows. In fish using both blue/danger- and SFPE-signals (Fig. 7e), the dEN is highly activated while the fish stays in the blue region during Go trials, due to the summation of the two signals. This activation can strongly destabilize the stay state and quickly induce a stay-to-move transition. Fig. 7e illustrates hypothetical potential landscapes showing how such transitions occur across behavioral states. Destabilization by the dEN flattens the potential landscape (orange line), facilitating transitions to the other state over the reduced barrier. Then, once the fish begins to move forward and receives scenery flow feedback, the SFPE disappears, and the discrete attractor-like potential landscape partially recovers, trapping the state in the move state (blue line). Behaviorally, this process would result in a short halt duration (Fig. 7f), as in the case of the real fish with both blue- and SFPE-coding activities in the Dc-indirect pathway-dEN circuit. In fish using blue/danger signal alone (Fig. 7g), the dEN modestly destabilizes the two behavioral states by the blue cue regardless of whether the fish is staying or moving (blue line). In this type of fish, the behavioral state would be equally stable at both stay and move states, resulting in longer halt duration (Fig. 7h), as in the case of the real fish with blue-coding activity alone in the Dc-indirect pathway-dEN circuit.

### Simulation of behavioral state transitions by pallidal output activity

To test whether the dEN can mediate such destabilization and state transition processes, we performed mathematical modeling (Fig. 8a). Note that the dEN neurons can provide uniform inhibitory output to all the components of the model Dc circuit. Simulation of this circuit with no dEN activity (as *dEN* = 0) from various initial values resulted in state transitions toward either (*Pmove*, *Pstay*) = (1, 0) or (0, 1) state, which promotes the behavioral move state or stay state, respectively (Fig. 8b, leftmost). However, when the dEN is highly activated (as *dEN* = 1.0), these two neural states were destabilized with flattening of the potential landscape, and two more stable points were added at (*Pmove*, *Pstay*) = (0, 0) and (1, 1) (Fig. 8b, rightmost). This is because the inhibitory output from the dEN abolished the mutual inhibition and replaced it with the uniform inhibition of the Pmove and Pstay neurons. Such destabilization occurred in a dEN activity level-dependent manner (Fig. 8b and Extended Data Fig. 9), and therefore, graded changes of dEN activity were converted into graded destabilization of the Dc activity state.

**Fig. 8:**
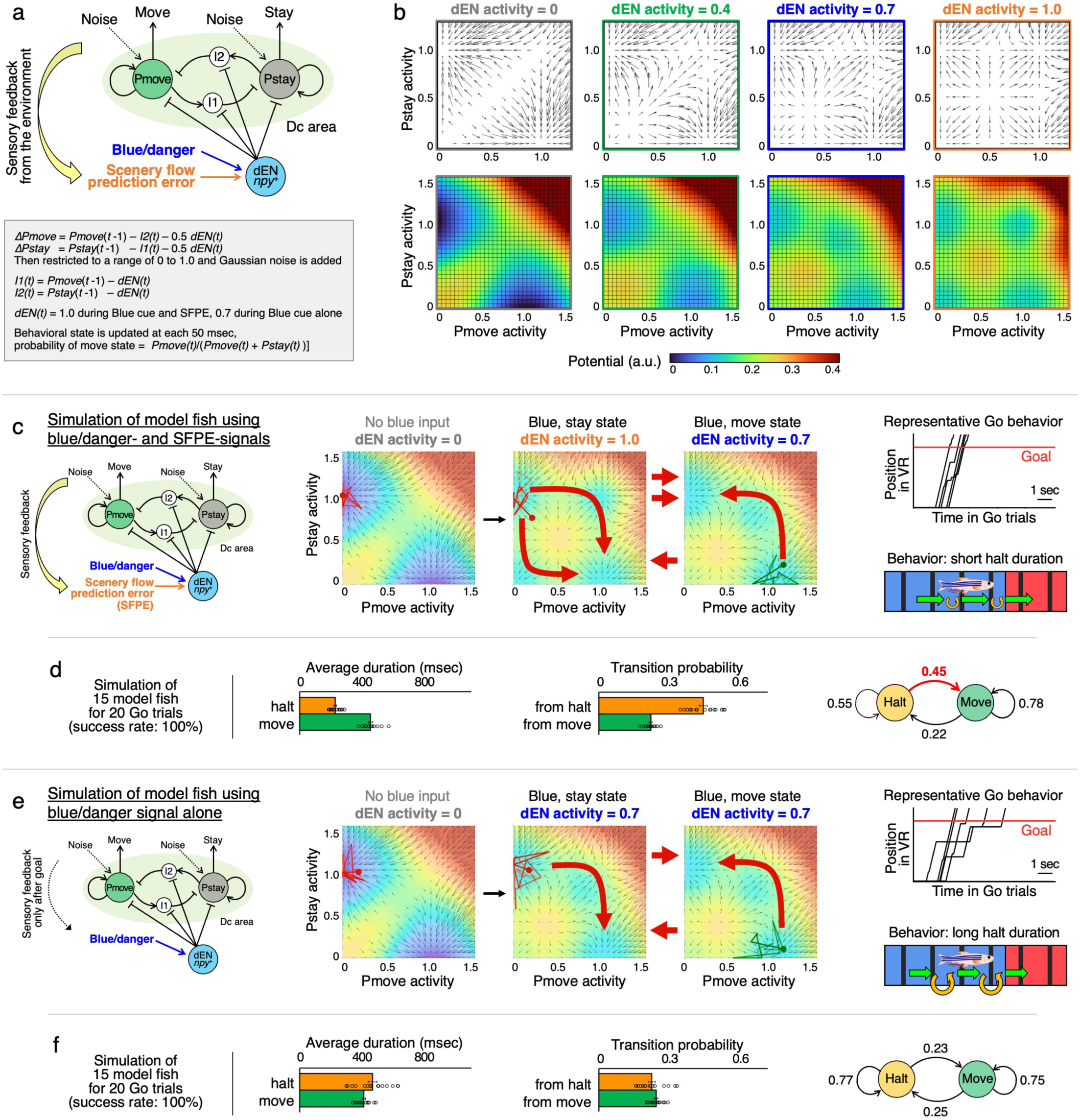
Mathematical modeling of neural and behavioral destabilization by the dEN activation during the Go/No-go task. **a**, (Top) Structure of the circuit model used in the simulations. (Bottom) Equations for simulating this circuit model. The I1 and I2 neurons indicate local inhibitory interneurons that mediate mutual inhibition between Pmove and Pstay neurons. See Methods for the details. **b,** (Top) Vector fields in the activity state space of Pmove neurons (*x*-axis) and Pstay neurons (*y*-axis) under different levels of dEN output activity. In this model, dEN activity ranges from 0 to 1.0. For Pmove and Pstay neurons, an activity value of 1.0 is the maximum value before adding Gaussian noise. Each arrow indicates a vector of state transition obtained by simulation of the model for one time step from the various initial values arrayed in the *x-y* coordinate. (Bottom) Color maps indicate potential landscapes calculated by the line integral of the corresponding vector field (see Methods for the details). State transitions tend to occur from a high potential state to a low potential state. **c,** Simulation of Go trials in model fish using blue/danger- and SFPE-signals. See also Supplementary Video 8. (Left) Schematics of this model. (Middle) Transition dynamics of neural activity states during escape behavior. Positions of dots in the state space indicate neural states (*Pmove*(*t*), *Pstay*(*t*)), and colors of dots indicate behavioral states (red, stay; green, move) at particular time points. Trailing lines connected to the dots indicate the recent history of the dots’ position. Red arrows indicate possible transition routes to the other state. (Right) Representative positional changes during five simulated Go trials, and an illustration of this behavioral pattern with shorter halt duration. **d,** (Left) Average halt and move durations in the model fish using blue/danger- and SFPE-signals (n = 15 model fish). The halt duration was shorter than the move duration, as observed in the real fish with blue/danger- and SFPE-coding Dc ensembles (see also Fig. 3c, top). (Middle) Transition probability during halts and moves in the same dataset. The transition probability during halts was higher than that during moves, as observed in the real fish (see also Fig. 3c, middle). No statistical tests were performed, as n is arbitrary in simulations. (Right) A state transition model between the halts and moves according to the middle panel. **e,** Same as in **c**, but for model fish using the blue/danger signal alone. See also Supplementary Video 9. (Right) This model resulted in the behavioral pattern with prolonged halt duration. **f**, Same as in **d**, but for the model fish using blue/danger signal alone (n = 15 model fish). These model fish showed similar levels of halt and move transition probabilities, as observed in the real fish with a blue/danger-coding Dc ensemble alone (see also Fig. 3d).

Simulation of the Go/No-go task by this model reproduced the behavior of real fish. To stochastically induce state transitions over time, we added constant Gaussian noise to the Pmove and Pstay neurons’ activity at each time step. In model fish using blue/danger- and SFPE-signals (Fig. 8c, Supplementary Video 8), the dEN was activated at the highest level while the fish stayed in the blue region during Go trials, and the flattening of the potential landscape opened two transition routes, one through (*Pmove*, *Pstay*) = (1, 1) and the other through (0, 0). This prompted a transition from the stay state to the move state, which abolished the SFPE and partially restored stability of the neural state by closing the route back to the stay state through (0, 0). Behaviorally, this process resulted in shorter halt duration and higher transition probability from halts than that from moves (Fig. 8d), as observed in real fish with blue/danger- and SFPE-coding Dc ensembles (see Fig. 3c). In contrast, in model fish using the blue/danger signal alone (Fig. 8e, Supplementary Video 9), the dEN was modestly activated and opened only one transition route throughout Go trials, resulting in halt duration and transition probability during halts similar to those during moves (Fig. 8f), as observed in real fish with a blue/danger-coding Dc ensemble alone (see Fig. 3d). In both models, the dEN output activity eventually guided behavioral state transitions toward the goal through destabilization of neural and behavioral states.

### Validation of the neural-state destabilization model using the experimental data

Lastly, we examined whether a transient absence of scenery flow feedback causes destabilization of the Pmove- and Pstay-neurons’ activity states in the real fish. By analyzing our previous data from Torigoe et al.,^26^ we compared their transition probability in desired [blue with scenery flow] and undesired [blue without scenery flow] time frames during Go success trials after learning. For this, we inferred firing patterns from the calcium traces by deconvolution with the decay time constant of G-CaMP7 (see Methods). Then, in each neuron, the estimated firing patterns were binarized into ON and OFF states by using the root mean square as a threshold, and the ON/OFF transition probabilities were calculated for [blue with flow] and [blue without flow] time frames separately. Distributions of this probability were plotted for each fish (Extended Data Fig. 10a). In the fish with shorter halts, this distribution was sparser along the [blue without flow] axis, due to less sampling of [blue without flow] time frames. To evaluate changes in transition probability across neurons, we linearly fit the distribution in each fish and measured the angle of inclination *θ* of the fitted line. The *θ* values higher than 45° were observed particularly in the fish with shorter halts, indicating biased elevation of transition probability in [blue without flow] than in [blue with flow] time frames in these fish (Extended Data Fig. 10b). This tendency was observed in both OFF-to-ON and ON-to-OFF transitions, and in both Pmove and Pstay neurons (Extended Data Fig. 10c, d), consistent with destabilization of ongoing neural activity states in our model. These results indicate that behavioral efficiency correlates with greater destabilization of neural states during no scenery flow time frames (undesired sensory feedback) rather than during scenery flow time frames (desired sensory feedback).

## Discussion

### The zebrafish pallidal output induces state destabilization

We have shown here that, in zebrafish, pallidal output neurons in the dEN uniformly broadcast the integrated negative value signal to the Dc. Optogenetic activation of this output pathway destabilized ongoing behavioral states and facilitated transitions to the other state. Based on these findings, we propose a neural circuit model in which dEN activity destabilizes Dc activity states, thereby guiding behavioral state transitions for proper goal-directed behavior. In this model, dEN activation flattens potential barriers, enabling exploration of other neural states through non-directional transitions. Once an appropriate behavioral state is reached, dEN activity is reduced, restoring the barrier to sustain that state. This process eventually leads fish to the behavioral goal, where dEN activity and the negative values are minimized.

Our model suggests that when a negative-value signal destabilizes ongoing behavioral states, it naturally guides behavioral state transitions toward reducing the negative value itself. In principle, this mechanism can also mediate behavioral choice among three or more options, which fish would encounter in their natural environment. This is because the dEN output can guide state transitions toward any Dc activity state that minimizes the output activity through random search by destabilization. This mechanism can explain how fish find an appropriate motor state under a given situation from many other motor attractor states. As for the motor output pathway from the Dc, zebrafish pallial projection neurons target various diencephalic and midbrain nuclei, including the tectum, although lacking the cortico(pallio)spinal tract.^21^

The dEN output was already active during successful Go trials in the initial phase of learning, although Go success trials were still infrequent at this phase (Extended Data Fig. 2b, c). This observation suggests that, even before blue-cue-dependent activation of the dEN is established, occasional dEN activation may facilitate escape behavior by destabilizing ongoing behavioral states, as a form of trial and error.

### Analogy of our model with simulated annealing in computer science

The destabilization mechanism proposed here resembles simulated annealing in computer science. This analogy has also been applied to the flattening of the brain’s energy landscape under psychedelics.^7^ In simulated annealing, a system is transiently heated so that it can escape a local minimum and explore other configurations, and then cooled so that it settles into a new minimum.^41,42^ In our model, dEN activation lowers the barriers between Dc activity states, which enhances the effect of heat and thus increases the fluctuation of the states. Once an appropriate state is reached, dEN activity declines, and the barriers are restored. Thus, the pallidal output may implement an annealing mechanism to search for appropriate behavioral states.

### A possible link to active inference

In our model, dEN activation destabilizes Dc activity and facilitates behavioral transitions toward minimizing dEN activity. Because the dEN integrates multiple negative value signals, including SFPE and blue/danger-coding signals, minimizing the dEN output would also minimize the SFPE, which encodes prediction error relative to the learned desired state (scenery flow). This process is conceptually similar to active inference, in which expected free energy is minimized through choosing actions that reduce the discrepancy between the current state and a desired sensory state (a preference prior).^30,31,43^ In addition, our dataset of Dc ensemble activities during learning has already provided experimental references for theoretical estimates of learning-dependent circuit reorganization based on active inference.^44^ Further theoretical study incorporating the current dataset will elucidate how the basal ganglia serve as a neural substrate for behavioral choice based on active inference.

Alternatively, the SFPE signal could be interpreted not as a prediction error but as a conjunctive danger signal in association with “blue without scenery flow.” Even under this interpretation, the SFPE signal must be computed by integrating the color cue and scenery flow information in some way (for example, the (ii) blue/flow-coding neurons in Fig. 6h). How such interaction occurs to generate the SFPE remains unknown, and further studies are required to elucidate the underlying circuit interactions. Although we did not address this issue in the present study, it does not affect our main conclusion: under either interpretation, destabilization of neural states that follows the temporal profile of this signal appropriately guides behavioral state transitions.

### Comparison between the fish and mammalian basal ganglia

Our observations may imply that the teleost pallio-basal ganglia circuit is evolutionarily related to the mammalian medial prefrontal-ventral striatal-ventral pallidal-mediodorsal thalamic circuit (mPFC-VS-VP-MD circuit, or the ventral loop). The ventral striatum and the ventral pallidum are thought to retain features of the ancestral cortico-basal ganglia circuit,^45^ which could also be shared with teleosts. The mammalian VS encodes value information for goal-directed behavior,^46^ and the VS indirect pathway neurons mediate inhibition of unsuccessful behavior.^47,48^ Moreover, mammalian VP inhibitory neurons also project to the mPFC in addition to the MD,^49,50^ as in the case of the fish dEN. Therefore, the mammalian ventral loop, particularly the mPFC-VS-VP-mPFC circuit, may support goal-directed behavior through an operational principle similar to that of the teleost basal ganglia.

### Putative involvement of the dopamine pathway in our model

We also found clear differences in the learning-dependent activities of the indirect and direct pathways of the fish striatum. In the indirect pathway, initially non-responsive neurons became responsive to the blue cue after learning. In addition, the SFPE-coding neurons also appeared after learning, particularly in fish with efficient escape behavior. Since the blue/danger- and SFPE-signals are prevalent Dc activities associated with punishment, these indirect-pathway activities may have been acquired through potentiation of the Dc-striatal indirect-pathway synapses by punishment. The zebrafish striatum also receives a projection from dopaminergic neurons.^51^ This raises the possibility that this potentiation is mediated by the D2 dopamine receptor, which detects dopamine dips during punishment and potentiates the cortico-striatal indirect-pathway synapses in mammals.^52,53^

In contrast, in the fish striatal direct pathway, initially tail-movement-coding neurons acquired responsiveness to the blue cue by learning. The acquisition of this activity may have been mediated by the D1 dopamine receptor, which detects dopamine release during successful outcome and potentiates cortico-striatal direct-pathway synapses that were active just before dopamine release in mammals.^52,54^ Potentiation of this pathway may also facilitate the initiation of the tail movement to avoid punishment, in coordination with the indirect pathway. Since we did not address the role of the direct pathway in this study, future studies will be needed to determine how it contributes to inducing learned behaviors.

### Zebrafish as a model animal to study the basal ganglia circuit

The teleost model of the pallio-basal ganglia circuit provides a versatile platform for investigating the mechanisms underlying basal ganglia function. For example, in adult zebrafish, the Dc and striatum are separated by approximately 400 µm along the dorsoventral axis. Therefore, it is technically feasible to semi-simultaneously image neural activity in the Dc and striatum during learning by rapidly alternating the focal planes.^55^ In addition, combining this approach with cell-targeted optogenetic stimulation^56^ would allow selective activation of functionally defined Dc neurons while simultaneously monitoring striatal responses. Such experiments will open new avenues for investigating a variety of intriguing phenomena, including striatal gating of cortical signals,^57^ dopamine-dependent plasticity,^52–54^ and learning-dependent changes in cortico-striatal functional connectivity. In addition, it has been recently reported that the zebrafish pallium is functionally regionalized with a hierarchical organization,^58^ as in the case of higher vertebrates. Together with recent advances in light-microscopy-based connectome analysis using expansion microscopy,^59^ the zebrafish telencephalon provides a promising platform for anatomical and functional connectomics to elucidate the circuit mechanisms of learning and decision-making.

## Methods

### Animals

All protocols were reviewed and approved by the Animal Care and Use Committees of the RIKEN Center for Brain Science (Protocol number: W2023–125, Approval number: 2024-014(1)). This study used adult zebrafish (RIKEN-Wako, Saitama, Japan) carrying the following transgenic alleles. The *Tg(npy:zGFF)^rw0164^*, the *Tg(UAS:eNpHR3.0-EYFP)^rw0165^*, and the *Tg(UAS:ChrimsonR-tdTomato)^rw0166^*lines were established in this study. The transgenic lines published previously and used in this study are as follows. The *Tg(penkb:zGFF)^rw0159^,* the *TgBAC(tac1:GAL4VP16)^rw0157^,*the *TgBAC(nkx2.1:GAL4VP16)^rw0160^,* the *TgBAC(npy:GAL4VP16)^rw0161b^*, and the *TgBAC(gad1b:GAL4VP16)^rw0162b^* are from Tanimoto et al.^21^ The *TgBAC(camk2a:GAL4VP16)^rw0154a^;TgBAC(vglut2a:Gal4)^nns20Tg^;Tg(UAS:G-CaMP7)^rw0155^*is from Torigoe et al. and Satou et al.^26,60^ The *Tg(UAS-hsp:zGCaMP6s)* is from Muto et al.^32^ The *Tg(UAS:GFP)^zf82^* is from Asakawa et al.^61^ The *Tg(UAS:DsRed2)^rw0135^*is from Agetsuma et al.^62^ The *Tg(UAS:AcGFP-2A-WGA)^rw0318Tg^*is from Takeuchi et al.^63^ The *TgBAC(gad1b:loxP-RFP-loxP-GFP)^nns26Tg^*is from Satou et al.^60^ The UAS:G-CaMP7 plasmid used for the *Tg(UAS:G-CaMP7)^rw0155^* line is from Yabuki et al.^64^ The *Tg(UAS:ChrimsonR-tdTomato)* line used in this study consists of two strains, the *Tg(UAS:ChrimsonR-tdTomato)^rw0166^*and the *Tg(5xUAS-hsp70l:ChrimsonR-tdTomato, myl7:mCherry)^nub119Tg^*, which is from Hagio et al.^35^ All the transgenic zebrafish were maintained in the nacre background^65^ and selected by their fluorescence patterns in the larval stage. Fish were maintained in 7-L tanks with continuous water exchange at 28.5°C under the 14-hour light/10-hour dark cycle, and male or female adult fish aged more than 3 months were used for the experiments.

### Generation of transgenic animals

For the generation of the *Tg(npy:zGFF)^rw0164^* knock-in transgenic line, the CRISPR/Cas9 knock-in strategy was used as described previously.^21,66^ The sgRNA sequence of TGAAGATGTGGATGAGCT, which targets the antisense strand starting 34 bp downstream of the start codon located in exon 2 of the *npy* gene, was used. The PAM sequence was GGG. Homology arms were designed as 482-bp and 627-bp flanking sequences of the upstream and downstream regions of the insertion site, respectively. The *hsp70* promoter, *zGFF* coding sequence,^61^ and poly-A tail were inserted at the cleavage site, located 28 bp downstream from the start codon of the *npy* gene.

For generation of the *Tg(UAS:eNpHR3.0-EYFP)^rw0165^* line, a plasmid construct pTol2-UAS-eNpHR3.0-EYFP was generated from pLenti-CaMKIIa-eNpHR 3.0-EYFP, which was kindly provided by Dr. Karl Deisseroth (currently available from Addgene, plasmid #26970; http://n2t.net/addgene:26970; RRID:Addgene_26970),^33^ by introducing an eNpHR 3.0-EYFP fragment into a pTol2-UAS vector. For the generation of the *Tg(UAS:ChrimsonR-tdTomato)^rw0166^*line, a plasmid construct pTol2-UAS-ChrimsonR-tdTomato was generated from pCAG-ChrimsonR-tdT, which was a gift from Edward Boyden (Addgene plasmid #59169; http://n2t.net/addgene:59169; RRID:Addgene_59169),^36^ by introducing a ChrimsonR-tdTomato fragment into a pTol2-UAS vector. For generation of the transgenic lines, these constructs were injected into the RIKEN-Wako strain using the *Tol2* transposon-mediated system for higher efficiency of transgenesis.^67^

### Immunohistochemistry

Immunohistochemistry was performed as described previously.^21^ Briefly, adult zebrafish brains fixed in 4% PFA in PBS (Nacalai 09154-85) were sliced into 75 µm vibratome coronal sections. All the primary and secondary antibodies used in this study are listed in Supplementary Table 1. All the sections were counterstained with DAPI (Invitrogen D1306). For permeabilization, PBS + 0.1% Triton X-100 was used throughout the primary and secondary antibody reactions. During the primary antibody treatment, 1% blocking reagent (Roche #11096176001) was also added to the buffer. The obtained sections were observed by a confocal laser scanning microscope (C2+, Nikon) with 5 µm or 2.5 µm intervals between confocal optical sections. To show the entire view of the section, maximum intensity projection (MaxIP) of all the optical sections from a 75-µm-thick section was used in Fig. 4a, leftmost panel; Fig. 4j, left panel; Extended Data Fig. 4b and 4c, leftmost panels; Extended Data Fig. 8a-b and 8c-d. For Extended Data Fig. 8e, MaxIP of 4 optical slices with 1.25 µm intervals was used. All the other images are single optical sections obtained by confocal imaging.

### Preparation of head-fixed adult zebrafish

To perform 2-photon imaging of the telencephalic neural activity, adult zebrafish were head-fixed under the objective lens, as described previously.^26^ Briefly, fish were anesthetized with 0.02% MS-222 (Nacalai 14805-82) and mounted on a hand-made surgery apparatus. During surgery, water with 0.02% MS-222 was continuously perfused to the mouth of the fish. The skin above the skull over the telencephalon and the tectum was removed using micro knives (10315-12, Fine Science Tools). After drying the surface, the dental bond (Scotchbond Universal Adhesive, 41255, 3I) was applied on the tectal skull and illuminated with blue LED light for 10 sec. A U-shaped stainless steel bar was attached to the tectal skull as a connector to the fixation apparatus with fluidic dental cement (Filtek Supreme Ultra, 6032XW, 3I) and illuminated with blue LED light for 10 sec. Then, the fish, the fixation apparatus, and a custom-made harness were assembled. After recovery from anesthesia by perfusion with water, the head-fixed fish and the assembly were transferred to the tank with the virtual reality environment. The fish was kept in the dark condition for ≥ 30 min under the microscope to allow it to habituate to the tethered situation with continuous perfusion of water.

### Installation of the cranial window

Motivated by the approach proposed in the previous study in mice,^37,38^ we have developed the following method. To acquire fluorescent signals brightly from the striatum located in the deeper part of the telencephalon, we installed cranial windows during the preparation of the head-fixed zebrafish. Before the installation of the cranial window, the fish were kept and adapted to teleost Ringer solution^24^ overnight. Then, all the following experimental procedures were performed in teleost Ringer solution instead of water. In addition, the dental bond was applied to both the tectal and telencephalic skull during the head fixation. Cranial windows were installed after the head fixation and before recovery from the anesthesia. We carefully cracked and removed the telencephalic skull using sharp forceps (Dumont #5SF, FST) and exposed the brain surface. Then, a piece of 12.5 µm-thick PFA Teflon microsheet (46850887, Yodogawa), which was cut to the approximate size of the fish head (2-3 mm width and 3-4 mm height), was placed so that it did not contain any bubbles between the sheet and the brain. Then, a gap between the sheet and surrounding skulls was carefully filled with the dental cement, which is connected to the tectal part of the cement. On top of the Teflon microsheet, a 3D-printed ring (∼2.0 mm in outer diameter and ∼0.1 mm in thickness) printed by Form3 (Formlabs) with Clear standard resin (Formlabs) was placed, and the sheet and the ring were embedded with dental cement to cover the surrounding area of the window. This protocol was applied to all the fish for imaging of direct-pathway neurons (n = 5 fish) and to a subset of fish for imaging of indirect-pathway neurons (n = 4 fish out of 8 fish).

### Go/No-go task in virtual reality

The VR experiment was performed essentially as we previously described,^26,68^ with the following modifications. The VR environment consisted of four LCDs (Good display, Model No. GD70MLXD), which presented the visual stimuli. For the three side panels, a commercially available privacy filter (EVF-HLPR12WN, Kokuyo) was used to prevent the display light from directly entering the objective lens. The virtual reality system was controlled by custom-written programs based on LabVIEW (National Instruments). The tail of the head-fixed fish was illuminated with infrared LED light (850 nm; KBG00000002, MeterMall), and was monitored by a USB camera (BSW200MBK, BUFFALO) with its IR cut filter removed. During this recording, a centerline lying along the body was continuously detected and analyzed to calculate the tail bending angle as described previously.^26^ The tail bend angles less than an angular threshold (mostly at 6 degrees, occasionally 5 degrees if the fish was small or moved subtly) were ignored and replaced with 0 degrees. The history of this tail-bending angle for the past 10 frames (*i.e.*, 1 sec) was used for calculation of tail beat frequency by Fourier transformation. This tail beat frequency was multiplied by the gain value and used as the forward locomotion speed of the fish. During the ITI or the open-loop experiment, this gain value was set to 0, *i.e.*, the fish could not move forward by tail movements, and no scenery flow feedback was delivered from the VR. During the Go or No-go trials, this gain value was set to 10, and fish could either move forward or stop by flipping or not flipping their tail. The behavioral video was captured and analyzed in real time at 20 Hz, and the data points were saved at 10 Hz.

The VR environment shown on the LCDs was created using Unity (Unity Technologies). Fish were located in a one-dimensional VR arena with a white, blue, or red background color with black stripes perpendicular to the direction of swimming. During ITI, the background color between the black stripes was white. During the Go trials, the background color of the near side of the fish in the VR arena was blue, and that of the far side was red. During the No-go trials, the background color of the near side of the fish was red, and that of the far side was blue. In this environment, fish were trained for the Go/No-go task, which consists of repetitions of 15 sec of ITI and 10 sec of trial. Before the training, fish were kept separated in individual 1 L tanks overnight. After the head fixation and habituation for more than 30 min in the dark, the adaptation session with 10 Go trials and 10 No-go trials in random order (20 trials in total) was initiated without the application of electric stimuli. After this adaptation session, the training sessions consisting of 30 Go trials and 30 No-go trials in random order (60 trials in total) were initiated, with inter-session intervals of 15–20 min without presentation of visual stimuli. In the training sessions, a weak and brief (5 V/cm for 1 sec) electric stimulus was applied as a punishment from needle electrodes near each side of their body at the end of each failure trial. Success or failure of each trial was determined by the fish’s position at the end of the trial; if the fish ended the trial in the blue region, it was a failure. If the fish ended the trial in the red region, it was a success. Each fish performed 2–10 training sessions, and the data obtained from learner fish were separated into the initial phase of learning, the intermediate phase of learning, and the after-learning phase according to Fig. 2b, bottom. If the initial phase contained fewer than three Go success trials, the earliest Go success trials from the intermediate phase were added to the initial phase to obtain three in total. A fish was defined as a learner when the success rates of the past 10 Go trials and past 10 No-go trials both exceeded 0.8. The open-loop experiments were performed after recording the after-learning phase in all the fish presented in this study. In some fish, rule reversal experiments were also performed after this open-loop experiment. During reversal learning sessions, the definition of success and failure was switched as follows: if the fish ended the trial in the red region, it was a failure. If the fish ended the trial in the blue region, it was a success.

### Classification of behavioral states

Behavioral states were classified into halt states and move states as previously described.^26^ The trajectories of the escape behavior (before reaching the goal) during Go success trials after learning were divided into time frames with positional change in the virtual arena (*pos*(*t*) – *pos*(*t*-1) > 0) and time frames without positional change (*pos*(*t*) – *pos*(*t*-1) = 0). The former time frames were classified as move states. The latter time frames were further subdivided into latency (the initial non-moving frames before any move state in a trial) and halt states (subsequent non-moving frames following a move state). The average halt duration of each fish was calculated as the mean duration of each halt episode by dividing [total time spent in halt states] by [total number of halt state terminations]. Similarly, the average move duration was calculated by dividing [total time spent in move states] by [total number of move state terminations]. Transition probabilities during halt states and move states (Fig. 3c, d and Fig. 8d, f, middle panels) were calculated as the reciprocal of the halt duration and the move duration, respectively, and then converted to the probability per 100 msec time frame. For Fig. 3 and Extended Data Fig. 1, our previous dataset from Torigoe et al.,^26^ where we recorded activity from the *camk2a*^+^ Dc excitatory neurons, was used. In this dataset, 9 fish had blue/danger- and SFPE-coding Dc ensembles, and 19 fish had a blue/danger-coding Dc ensemble alone.

For the duration distributions (Extended Data Fig. 1c, d), halt and move events were pooled across fish within each group, and the transition probability per 100 msec (*p*) was calculated as the number of state transitions divided by the total number of 100-msec bins spent in that state, as described above. Then, the estimated time constants shown in these panels were calculated as *τ* = −100 msec/ln(1 − *p*). Moves truncated by goal arrival were included in the total duration but not counted as transitions from moves.

### Two-photon imaging

To visualize calcium signals, we used a two-photon microscope (A1R-MP, Nikon) with a water-immersion objective lens (25x Nikon CFI APO LWD Objective, 380 - 1050 nm, 1.10 NA, 2.0 mm WD, Nikon) and a Ti:sapphire laser (Mai Tai eHP DeepSee, SpectraPhysics) at a wavelength of 920 nm. Fluorescence emissions were collected using a GaAsP photomultiplier tube. Scanning was performed using a resonant scanner by unidirectional scanning, with 65 µsec per line and another 65 µsec per flyback time. The backlight of the LCDs of the VR system was turned on only during the first 15 µsec of each flyback time by TTL control (Master 9, AMPI). The imaged field was 509 × 509 µm (512 × 512 pixels). Imaging was performed using a piezo actuator (Nano-Drive, MCL) to observe three different focal planes separated by 16 µm for imaging of dEN axons in the pallium, and by 24 µm for imaging of striatal neurons. This resulted in approximately 300 msec per volume (including three frame acquisitions and piezo transfers), and an acquisition rate of ∼ 3.23 volumes per second. For recording of the striatum and the dEN axons, the virtual reality display at the bottom of the tank, which is dispensable for learning the task, was not used to improve signal-to-background ratio. Part of the fish samples for the recordings of indirect-pathway neurons (n = 2 fish out of 8 fish) and for the recordings of dEN axon termini (n = 2 fish out of 5 fish) were obtained using a different two-photon microscope system LSM710 (Zeiss), of which methodological details were already described previously.^26^

### Image processing

Image processing of the calcium imaging data was performed essentially as described previously.^26^ Image data analysis was performed using scripts written in ImageJ (ImageJ, NIH), LabVIEW (National Instruments), and MATLAB (MathWorks). The imaging data derived from a single slice were collected using the custom macro of ImageJ (NIH). The collected image data of each slice were processed to remove the displacement of the x–y axis during the experiment by two-dimensional Fourier transform to determine the displacement of individual images compared to the reference image, generated by temporal averaging of 1000 images with minimal displacement from the dataset. The distance of displacement in the frequency domain was calculated in the number of pixels along the x and y axes using cross-correlation, and each image was corrected for the shift in the x and y axes. After displacement correction, the images were then used to generate a standard deviation map (STD map), calculated as a map of standard deviation across all the time frames of the imaging in each pixel.

The ROI detection and analysis of pallial excitatory neurons was performed in the previous study.^26^ For the ROI detection of the dEN axon termini in the pallium, we used a slightly modified method from the one from Torigoe et al.^26^ To detect the ROIs corresponding to individual axon termini, we used the STD map. The axonal activity caused fluorescent intensity changes with greater deviation over time, resulting in higher STD values in the pixels inside the axons than in the pixels outside the axons. Then, we ranked the pixels by their STD values, and we took the 1st ranked pixel to calculate the correlation of intensity changes over time between this pixel and the surrounding pixels (51 × 51 pixels). We set a threshold value of correlation empirically for each fish, and if the correlation between this pixel and one neighboring pixel was higher than the threshold, that neighboring pixel was considered to be in the same ROI as this pixel. Using this process, a boundary of an ROI could be obtained. We repeated this process for the second-ranked pixel in the updated list to identify another ROI. This process was repeated for many ranked pixels in descending order until a threshold was reached (typically until a mean STD value among all pixels). After the detection of potential ROIs using this method, we checked the spatial overlap between the ROIs. If two ROIs overlapped, the ROI with the smaller area was removed. In addition, ROIs detected in the area of the image where there was apparently no brain tissue were also removed. For analysis of the neuronal activity in the striatum, cell ROIs were detected manually from the STD map as a round-shaped brighter spot compared to the surrounding background with a size of 10 to 14 µm in diameter, depending on the x-y resolution of the imaging.

After detecting ROIs, the fluorescent intensity of all the pixels belonging to each ROI was averaged in each time frame and obtained as temporal fluorescent intensity changes. We also calculated the background intensity by averaging all the pixels belonging to the background ROI in each time frame. The background ROI was defined as pixels below a threshold value in the STD map of each focal plane. The corresponding background intensity was subtracted from the temporal fluorescent intensity change of each ROI at each time point to calculate F(t), which was later used to calculate ΔF/F_0_. To match the F(t) data obtained at 3.23 volumes per second (for 3 focal planes) to the behavior data obtained at 10 Hz, we interpolated the F(t) data to generate F(t) traces at 9.69 Hz, and assigned one F(t) value to each time frame of behavior data based on the frame number of the corresponding fluorescent images saved during the experiment, as described previously.^26^ For the analysis of overall dEN axon activity (Fig. 4e, 5g, h, Extended Data Figs. 2c and 3b), a single large ROI covering the dEN axon-innervated area of the Dc was drawn in each of the three focal planes, and F(t) was obtained by summing the background-subtracted fluorescence intensities of all the pixels within the three ROIs, so that each focal plane contributed in proportion to its ROI area. F_0_ was defined at each trial by averaging F(t) during the last 5 sec of each ITI.

### Classification of striatal neurons

Striatal neurons were classified based on the correlation coefficient between the recorded neuronal activity and the simulated activity templates (Fig. 6g). The simulated activity templates were generated from the real sensory and behavioral data of each real fish. For this, temporal patterns of presence or absence of blue cue, tail movement (> 6 deg), and scenery flow were converted to square-wave patterns [Blue], [Flow], and [Move], which were set to 1 when present and to 0 when absent. Then, a new waveform [Blue] OR [Flow] was synthesized by logical OR operation of the square-wave patterns of [Blue] and [Flow]. A new waveform [Blue] OR [Move] was synthesized by logical OR operation of the square-wave patterns of [Blue] and [Move]. A new waveform [SFPE] was synthesized by logical AND operation of the square-wave patterns of [Blue] and (1−[Flow]). These synthesized square-wave patterns were convolved with an exponential decay kernel approximating the kinetics of GCaMP6s (*τ*_off_ = 1 sec)^69^ to generate neural activity-like patterns of the simulated activity templates (see Fig. 6g).

These simulated activity templates were then used to calculate Pearson correlation coefficients with the activity patterns of all the striatal neurons. For this calculation, 15 sec just after receiving the mild electric stimulus were omitted to avoid spurious correlation elicited by the punishment. Then, the activity patterns of each striatal neuron were classified into (control) “simple blue-coding neurons” if the correlation coefficients *r* with [Blue] template before and after learning are both ≥ 0.2; into (i) “blue/danger-coding neurons” if *r* with [Blue], [Flow], and [Move] templates before learning are all < 0.2 and *r* with [Blue] template after learning is ≥ 0.2; into (ii) “blue/flow-coding neurons” if *r* with [Flow] before learning is ≥ 0.2 and *r* with [Blue] OR [Flow] after learning is ≥ 0.2; into (iii) “blue/move-coding neurons” if *r* with [Move] before learning is ≥ 0.2 and *r* with [Blue] OR [Move] after learning is ≥ 0.2; into (iv) “SFPE-coding neurons” if *r* with [SFPE] template after learning is ≥ 0.2 (no restriction before learning due to heterogeneous activity patterns of SFPE-coding Dc ensembles before learning^26^). If one striatal neuron was classified into two or more response types, only the response type with the highest *r* after learning was adopted. The distributions of the classified *penkb*^+^ and *tac1*^+^ neurons’ *r* after learning into the five response types are shown in Extended Data Fig. 7a, b.

### Classification of Pmove and Pstay neurons

Pmove and Pstay neurons were detected from our previous dataset from Torigoe et al.,^26^ where we recorded activity from the *camk2a*^+^ Dc excitatory neurons using *TgBAC(camk2a:GAL4VP16);TgBAC(vglut2a:Gal4);Tg(UAS:G-CaMP7)* fish. Of the 9 fish with blue/danger- and SFPE-coding Dc ensembles and the 19 fish with a blue/danger-coding Dc ensemble alone in this dataset, we used 8 of the former and 16 of the latter for this analysis, because other fish were recorded with a different imaging condition to capture deeper regions of the dorsal pallium. For the detection of Pmove and Pstay neurons, ΔF/F_0_ traces were binarized into ON and OFF states by thresholding with the root mean square of the full recording trace in each neuron. Then, the temporal probability of move states (tail angle > 6 deg) was calculated for the time frames of activity ON states. If the calculated move probability during ON states was 1.5-fold higher than that during the entire time-course, the neuron was determined as a Pmove neuron. If the calculated move probability during ON states was lower than that during the entire time-course, the neuron was determined as a Pstay neuron.

### Optogenetics

For optogenetic manipulation of dEN axons in the pallium, we used a custom-made optical path additionally installed on the side of the A1R-MP (Nikon) two-photon microscope system so that it can deliver wide-field illumination of 561 nm light onto the entire view-field. The 561 nm single-photon light produced from LED M595L4 (Thorlabs) and controller DC2200 (Thorlabs) through bandpass filter FBH560-10 (Thorlabs), and collimating lens ACL2520U-A (Thorlabs) was used for both the optogenetic suppression and activation experiments.

For the optogenetic suppression experiment of dEN axons using *Tg(npy:zGFF)^rw0164^;Tg(UAS:eNpHR3.0-EYFP)^rw0165^*fish (Fig. 4i-m), 16x objective lens CFI75 LWD 16x W (Nikon), which allows wide-field light illumination to an area with approximately 1300 µm diameter, was used for the suppression of the bilateral pallium. For the adjustment of the relative position of the lens and fish, we imaged the EYFP signal with the A1R-MP and set the focal plane at a similar depth to that in the dEN axon imaging in the pallium (see Fig. 4b). Then, the midline was set at the center of the view-field, and the caudal end of the pallium at the edge of the view-field. The light illumination of 0.45 mW/mm^2^ was pulsed at 40 Hz with a 2 msec pulse duration. Illumination was performed during the Go or No-go trial (10 sec) after learning the Go/No-go task. To test the immediate effect of the optogenetic suppression, we alternated light ON and OFF blocks every 5 trials and repeated this for one training session with 30 Go and 30 No-go trials in random order (60 trials in total, with 6 ON and 6 OFF blocks). Then, we aggregated the trials for light ON and OFF phases for further statistical analysis. Control fish were non-Tg fish of similar age in the same genetic background (RW strain, nacre).

For the optogenetic activation experiment of dEN axons (Fig. 5a-e), triple transgenic lines of *Tg(npy:zGFF)^rw0164^;Tg(UAS:ChrimsonR-tdTomato)^rw0166^;Tg(UAS-hsp:zGCaMP6s)* fish and *Tg(npy:zGFF)^rw0164^;Tg(5xUAS-hsp70l:ChrimsonR-tdTomato, myl7:mCherry)^nub119Tg^;Tg(UAS-hsp:zGCaMP6s)* fish were used.^35^ As the control fish of this experiment, *Tg(npy:zGFF)^rw0164^;Tg(UAS-hsp:zGCaMP6s)* sibling or half-sibling fish were used. The light illumination condition was modified for simultaneous two-photon imaging of the dEN axons as follows. For delivery of the whole-field 561 nm LED light pulsed at 40 Hz with a 2 msec pulse duration, a 25x objective lens CFI APO LWD (Nikon), which allows illumination to an approximately 800 µm diameter, was used to unilaterally illuminate the light in the pallial Dc area with 1.25 mW/mm^2^ intensity. For simultaneous two-photon imaging during this activation experiment, we also used a dichroic mirror 488/561 VIS/IR (Nikon), which passes the 561-nm light and 920-nm two-photon laser while reflecting fluorescence at 500-550 nm. In addition, the 561-nm light was turned OFF during the unidirectional scanning of the two-photon imaging and turned ON only during the flyback time (65 µsec). This experiment was performed using naïve fish in a continuous white background color without any pre-training. The light illumination was applied as repetitions of 15 sec OFF phases and 10 sec ON phases for 20 trials. The recorded neural and behavioral responses were trial-averaged for these 20 trials and used for Fig. 5c, e. In this behavioral analysis of the control fish and the ChrimsonR^+^ fish, we did not consider short periods (< 1 sec) with the tail flip less than 6 degrees as a stay period when they occur during tail flipping, to avoid mis-fragmentation of the successive tail movement of the naïve control fish, which moved slowly and less robustly compared with the learner fish of the Go/No-go task.

### Evaluation of neural state stability in [blue with flow] and [blue without flow] time frames

Probabilities of ON/OFF state transitions of Pmove and Pstay neurons in [blue with flow] and [blue without flow] time frames (Extended Data Fig. 10) were calculated as follows. From the dataset of imaging of *camk2a*^+^ Dc excitatory neurons (original data from Torigoe et al.^26^), time frames during escape behavior in Go success trials after learning were extracted. According to the sensory state of each frame, these frames were divided into [blue with flow] and [blue without flow] time frames. Then, the corresponding calcium responses of *camk2a*^+^ Dc excitatory neurons were converted from ΔF/F_0_ traces to estimated firing response *r*_est_(*t*) traces by deconvolution. Assuming that the ΔF/F_0_ traces *c*(*t*) reflect the firing response convolved with an exponential decay kernel with the decay time constant *τ* of the calcium indicator, *r*_est_(*t*) was estimated as *r*_est_(*t*) = *c*(*t*) + *τ*·d*c*(*t*)/d*t*, where d*c*(*t*)/d*t* was approximated by the central difference [*c*(*t* + Δ*t*) − *c*(*t* − Δ*t*)]/(2Δ*t*) with Δ*t* = 100 msec (one time frame), and *τ* was set to the decay time constant of G-CaMP7 (*τ*_off_ = 470 msec).^70^ Then, the *r*_est_(*t*) traces were binarized into activity ON and OFF states by thresholding with the root mean square of the *r*_est_(*t*) traces in each neuron, and OFF-to-ON and ON-to-OFF transition events were detected for [blue with flow] and [blue without flow] time frames separately. For these [blue with flow] and [blue without flow] time frames, the probabilities of the OFF-to-ON and ON-to-OFF transition events (per 100 msec time frame) were calculated by dividing the counted number of transition events by the total time length of the collected time frames of [blue with flow] or [blue without flow] in each neuron. Then, these probabilities were compared between the [blue with flow] and [blue without flow] time frames by plotting each neuron’s calculated probability values on an X-Y plane for each fish. The plotted values of Pmove or Pstay neurons were then linearly fitted by the total least squares method (as *y* = *ax* + *b*) to calculate their angle of inclination *θ* = arctan(*a*) for each fish. We detected biased elevation of transition probability in [blue without flow] time frames as *θ* > 45°, and in [blue with flow] time frames as *θ* < 45°. Then, the dependency between this *θ* and the average halt duration was plotted for individual fish and fitted by a power approximation (as *y* = *ax^b^)*. The negative *b* values observed for both OFF-to-ON and ON-to-OFF transitions, and both for Pmove and Pstay neurons, indicate that shorter halt duration correlates with biased elevation of transition probability in [blue without flow] time frames.

### Mathematical modeling of the neural circuit

The neural circuit model was formulated in the form of population activities of Pmove and Pstay ensembles as continuous variables according to the equations as follows (Fig. 8a):

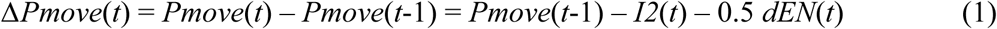

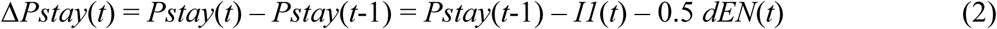

Therefore:

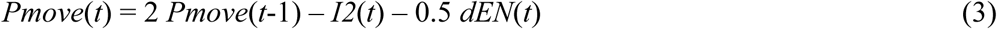

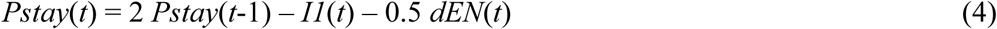

where *I1*(*t*) and *I2*(*t*) are:

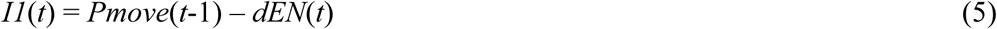

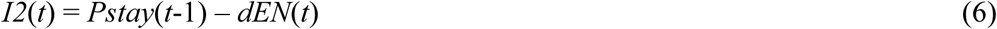

Note that after calculating *I1*(*t*) and *I2*(*t*), these values less than 0 were rounded to 0. After calculating *Pmove*(*t*) and *Pstay*(*t*), these values less than 0 were rounded to 0, and these values greater than 1.0 were rounded to 1.0.

The vector field plots shown in Fig. 8b were generated using the quiver function implemented in MATLAB (MathWorks). For each plot, the model was simulated for a single time step from a set of systematically varied initial values as (*Pmove*(0), *Pstay*(0)) = (*X*, *Y*), which are arranged in a grid parallel to the horizontal and vertical axes of the neural state space of *Pmove*(*t*) and *Pstay*(*t*), respectively. The difference between (*X*, *Y*) and the position at the next time step (*Pmove*(1), *Pstay*(1)) was calculated as vectors (*U*, *V*) under given constant dEN activity levels *dEN*_const_ = 0, 0.4, 0.7, or 1.0 as follows:

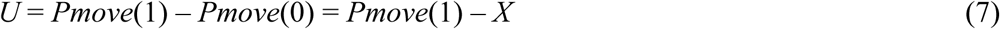

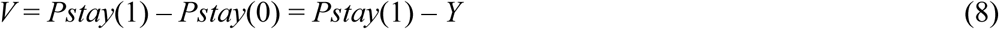

where *Pmove*(1) and *Pstay*(1) are:

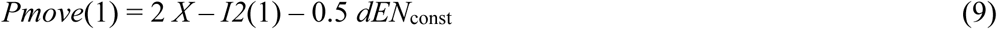

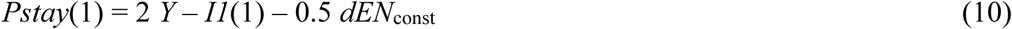

where *I1*(1) and *I2*(1) are as follows:

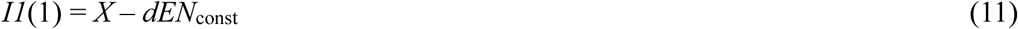

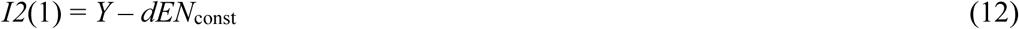

After calculating *I1*(1) and *I2*(1), these values less than 0 were rounded to 0. After calculating *Pmove*(1) and *Pstay*(1), these values less than 0 were rounded to 0, and these values greater than 1.0 were rounded to 1.0. The resulting values (*U*, *V*) are the displacement vectors originating from each (*X*, *Y*) point in the neural state space. These values were then used as input to the quiver function to visualize the local flow of the system in the neural state space. The resulting vector fields illustrate the direction and magnitude of attractor dynamics across the examined region of the state space. The distributions of potential landscape were subsequently estimated by averaging two potential distributions, which were calculated by the line integral of the obtained vector fields, one integrated from (*X*, *Y*) = (0, 0) to (1.5, 0) and then (*N*, 0) to (*N*, 1.5) for *N* = 0 to 1.5, the other integrated from (*X*, *Y*) = (0, 0) to (0, 1.5) and then (0, *N*) to (1.5, *N*) for *N* = 0 to 1.5. Then, the averaged potential distribution was visualized as an offset relative to its peak value (within the area of 0 ≤ *X* ≤ 0.5 and 0 ≤ *Y* ≤ 0.5), which was aligned with 0.25 in the color scale. Exact scalar potentials can be derived when the vector field is conservative, as in the case of *dEN*_const_ = 0. When the field is near-conservative, as in the case where *dEN*_const_ is higher than 0, approximate potentials can be constructed by the line integral.

Based on this circuit model, the Go/No-go task was simulated to reproduce the escape behavior of the real fish after learning. Sensory input affected the *dEN*(*t*) as follows: *dEN*(*t*) was 1.0 when there were both the blue cue and SFPE, 0.7 when there was only the blue cue, and 0 when there was the white cue. In Go trials, the blue cue was present until the fish reached the goal. In the simulation of Fig. 8c, SFPE was present whenever the simulated fish were not seeing scenery flow in the presence of the blue cue. To reproduce the behavioral response latency observed in the real fish, *dEN*(*t*) was set to 0 for the first 1500 msec of the trials as a waiting time in all the conditions. The other initial values were determined for the simulation of Go/No-go trials as follows:

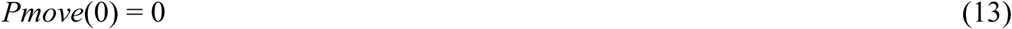

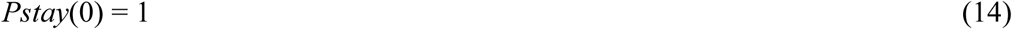

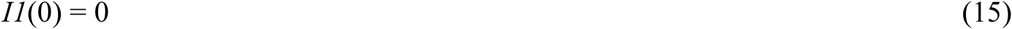

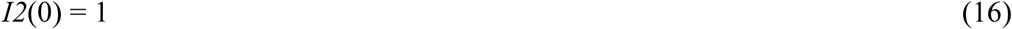

In the later time steps, temporal changes in the neural activity were calculated according to equations (1) to (6). After calculating *I1*(*t*) and *I2*(*t*), these values less than 0 were rounded to 0. Here, after calculating *Pmove*(*t*) and *Pstay*(*t*), these values less than 0 were rounded to 0, and these values greater than 1.0 were rounded to 1.0, and then Gaussian noise (mean = 0, standard deviation = 0.145) was added. Therefore, *Pmove*(*t*) and *Pstay*(*t*) tended to range from 0 to 1.0 and occasionally became higher than 1.0 due to the addition of the Gaussian noise. One trial was simulated for 10 sec with 0.1-msec time steps, and the behavioral states were updated with 50 msec time steps based on the *Pmove*(*t*) and *Pstay*(*t*) values of that time point. At each 50 msec, if *Pmove*(*t*) was less than 0.5 or *Pstay*(*t*) was greater than 0.5, the behavioral decision *D*(*t*) was 0, *i.e.*, the stay state. If *Pmove*(*t*) was greater than or equal to 0.5 and *Pstay*(*t*) was less than or equal to 0.5, *D*(*t*) was randomly sampled from a Bernoulli distribution Ber(*p*) according to the following probability *p* using the mnrnd function in MATLAB:

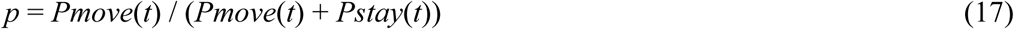

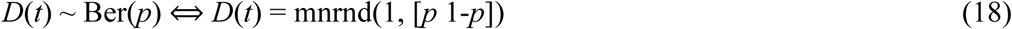

Positions of the fish *pos*(*t*) were initialized and updated each frame as follows:

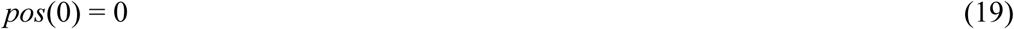

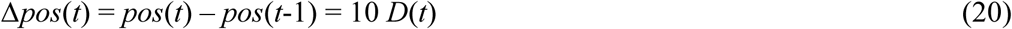

For each determination of *D*(*t*) as 1, *i.e.*, move state, the state lasted at least 50 msec until the next update. This caused an increase of *pos*(*t*) by 5000. The goal position *pos*_g_ was set to 100000. Therefore, for each Go trial, moving forward 1 sec in total (20 determinations of *D*(*t*) as 1 in total) led the simulated fish to the goal. For the statistical analysis shown in Fig. 8d and f, 40 trials (20 Go trials and 20 No-go trials) were simulated for 15 fish. Average halt duration was calculated for each fish across all the halt events during the Go success trials, as in the case of real fish from Torigoe et al.^26^

### Statistical Details

Two-tailed paired *t*-tests were performed using LabVIEW 2012 (National Instruments) or Excel for Mac ver 16.106 (Microsoft). Two-tailed unpaired *t*-tests were performed using Excel for Mac ver 16.106 (Microsoft). One-way ANOVA with Tukey’s HSD post hoc tests and Dunnett’s tests were performed using R (The R Project). The significance level *α* was set at 0.05 for all the statistical tests. Asterisks indicate levels of statistical significance as follows: * for *P-*value from 0.05 to 0.01, ** for *P*-value from 0.01 to 0.001, and *** for *P*-value less than 0.001.

## Data and Code Availability

All the data points, sample size, and p-values used to generate the figures will be provided as a supplementary spreadsheet file before acceptance of this manuscript. The calcium imaging and behavioral recording data used in this study will be deposited at Zenodo and made publicly available before acceptance of this manuscript. The MATLAB scripts and data used for the mathematical simulation will be deposited at Zenodo and made publicly available before acceptance of this manuscript. Raw microscopy data used in this paper will be shared by the corresponding author upon request. Any additional information required to reanalyze the data reported in this paper is available from the corresponding author upon request. All the transgenic lines and plasmids generated in this study are available from the corresponding author upon request.

## Acknowledgments

We thank Toshio Ohshima and Fumi Kubo for their institutional support, Peter Dayan for critical reading of the manuscript and insightful comments, Sten Grillner, Yu Mu, Kenji Doya, Jun Igarashi, and Ken-ichi Inoue for their discussion and comments, Karl Deisseroth and Ryunosuke Amo for providing the pLenti-CaMKIIa-eNpHR 3.0-EYFP plasmid, Edward Boyden for providing the pCAG-ChrimsonR-tdT plasmid, Masahiko Hibi and Wataru Koyama for providing the *Tg(5xUAS-hsp70l:ChrimsonR-tdTomato, myl7:mCherry)* line, Shin-ichi Higashijima for providing the *TgBAC(gad1b:loxP-RFP-loxP-GFP)* line, Yoshihiro Yoshihara for providing the *Tg(UAS:AcGFP-2A-WGA)* line and the G-CaMP7 plasmid, Junichi Nakai and Tetsuya Koide for providing the G-CaMP7 plasmid, Nobuhiko Miyasaka for providing the *Tg(UAS:AcGFP-2A-WGA)* line, Koichi Kawakami for providing the *Tg(UAS:GFP)* line, the *Tg(UAS-hsp:zGCaMP6s)* line, the *zGFF* construct, and the *Tol2* constructs, Yoshiko Takahashi for providing the *Tg(UAS-hsp:zGCaMP6s)* line, and Hisato Maruoka and Toshihiko Hosoya for their experimental support. We thank all the members of the Okamoto and Kubo laboratories for their support and advice, especially Yoshie Ito for experimental assistance. We thank the Research Resource Division of RIKEN CBS for fish care and technical assistance, and the National BioResource Project of Japan for providing zebrafish strains.

## Author information

### Contributions

H.O. conceived and supervised the project. Y.T., M.T., T. Islam, and H.O. designed the experiments and analyses. Y.T., M.T., and T. Islam built the virtual reality and computerized control and data acquisition systems. M.L. initiated the development of the cranial window protocol, and Y.T. refined it. Y.T. carried out the experiments and analyses. Y.T., H.K., and T.S. established transgenic lines. Y.T. and H.O. conceived the circuit model. Y.T. and T. Isomura designed the mathematical modeling. Y.T. performed mathematical modeling with the help of T. Isomura. Y.T. and H.O. wrote the manuscript.

## Extended Data Figure titles and legends

**Extended Data Fig. 1:**
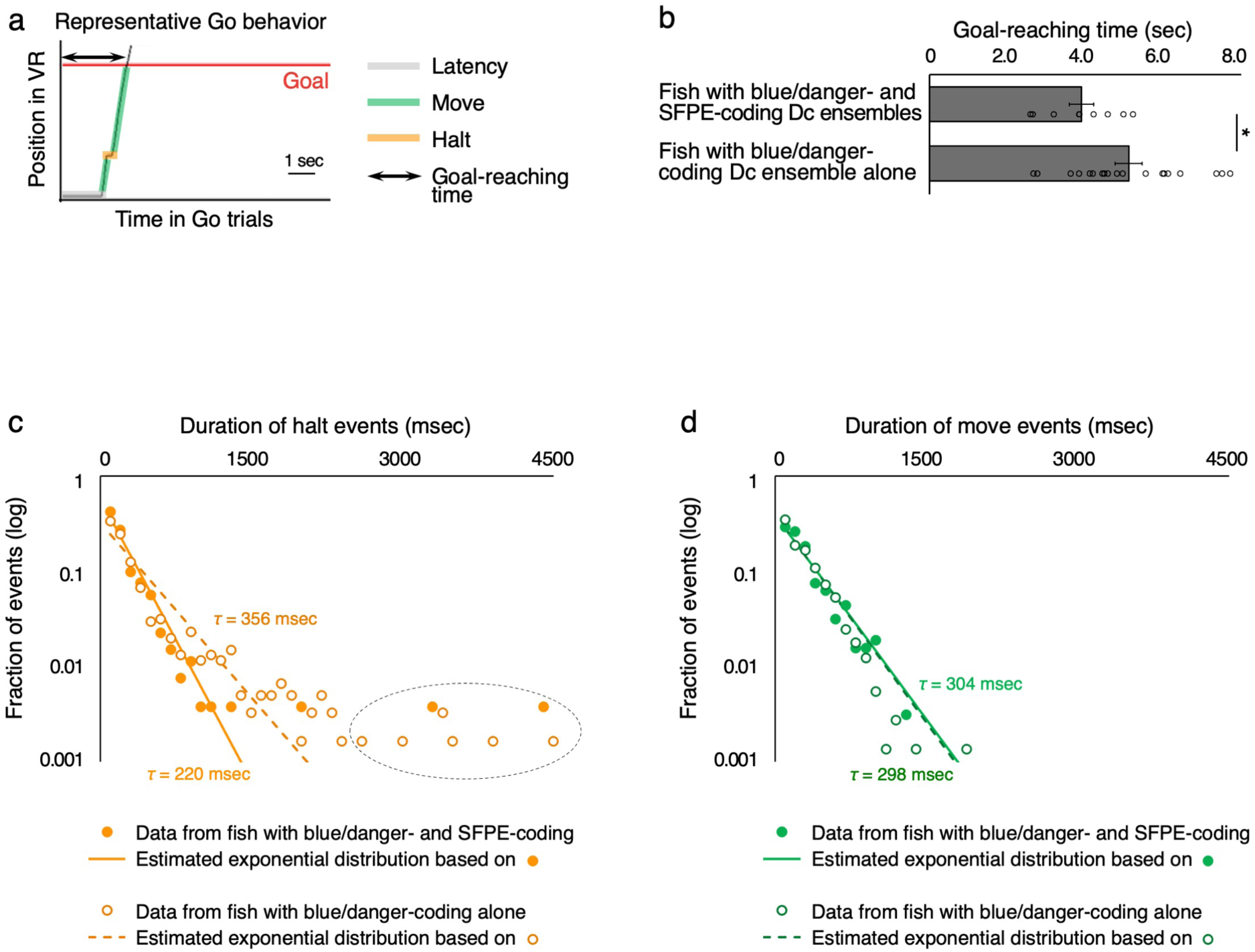
Different behavioral performances of the two types of learner fish. **a**, Illustration of the definition of the halt and move states during the learned escape behavior, overlaid on a representative Go success trial after learning. **b,** The bar graph shows mean ± s.e.m. of goal-reaching time of Go trials from the two groups of learner fish (original data from Torigoe et al.^26^). The fish with blue/danger- and SFPE-coding Dc ensembles (n = 9 fish) showed reduced goal-reaching time compared with fish with a blue/danger-coding Dc ensemble alone (n = 19 fish); \**P* = 0.036, two-tailed unpaired *t*-test. **c,** Distributions of halt durations in Go success trials after learning, pooled across fish with blue/danger- and SFPE-coding Dc ensembles (filled circles; n = 9 fish; 265 halts) and fish with a blue/danger-coding Dc ensemble alone (open circles; n = 19 fish; 606 halts). Each point shows the fraction of halts in a 100-msec bin, plotted on a logarithmic scale. Lines show the exponential distributions estimated from the observed data, whose time constants were calculated as *τ* = −100 msec/ln(1 − *p*), where *p* is the transition probability per 100 msec derived from pooled events across fish (see Methods). The dotted circle indicates rare occurrence of extremely long halt events that deviate from the exponential distribution. **d,** Same as in **c**, but for move durations pooled across fish with blue/danger- and SFPE-coding Dc ensembles (filled circles; n = 9 fish; 319 moves) and fish with a blue/danger-coding Dc ensemble alone (open circles; n = 19 fish; 732 moves). Note that move durations include moves truncated by goal arrival, which shortens the observed distribution compared with the estimated distribution.

**Extended Data Fig. 2:**
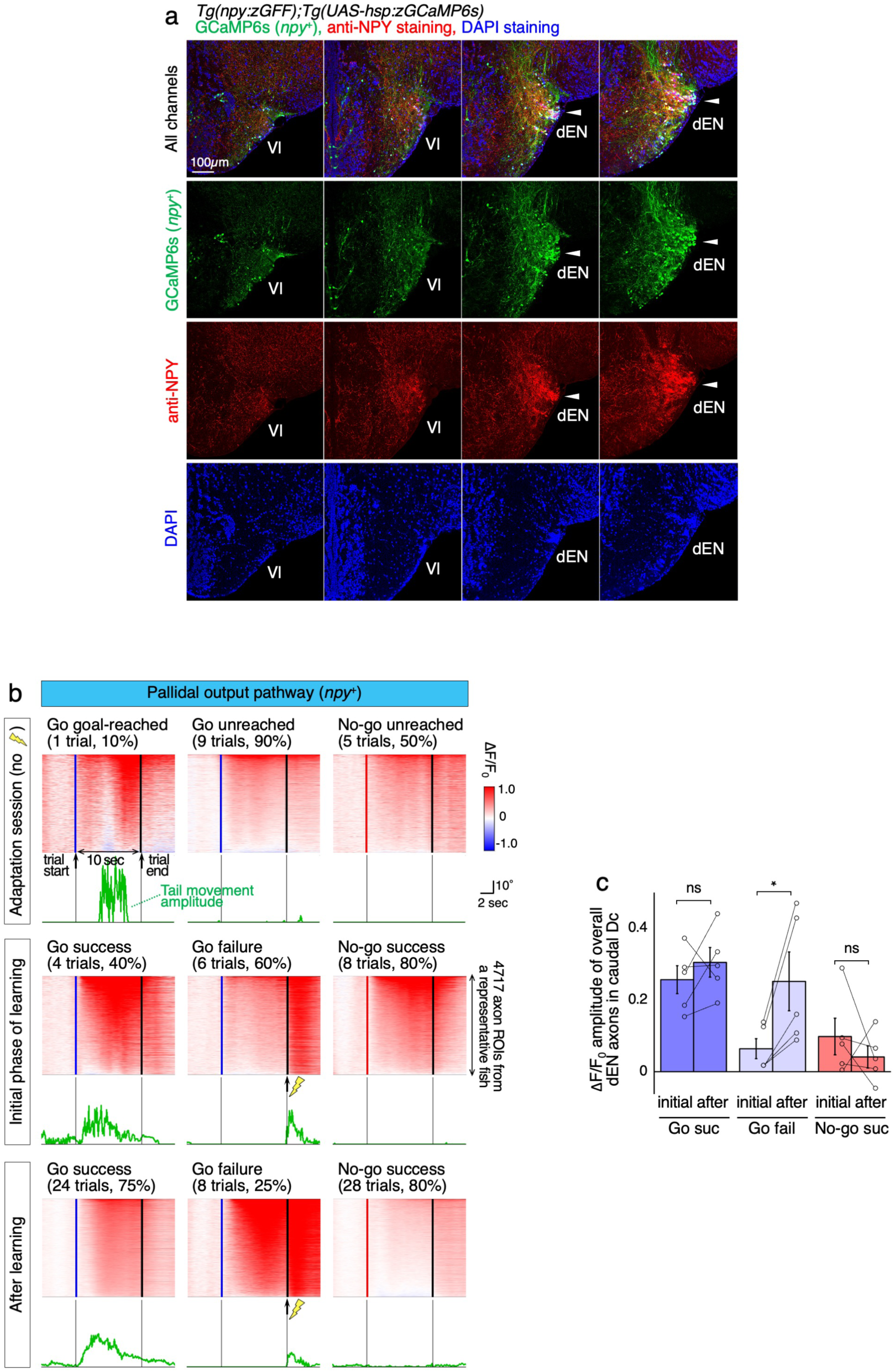
Learning-dependent activity changes of *npy*^+^ dEN axons in the Dc during Go success, Go failure, and No-go success trials. **a**, Immunohistochemistry for GCaMP6s (green) and NPY (red) expressed in *npy*^+^ dEN output neurons. Four successive coronal sections from the anterior (left panels) to the posterior (right panels) parts of the telencephalon are shown. The anterior two sections contain the NPY⁻ Vl region, and the posterior two sections contain the NPY^+^ dEN region. Blue channel indicates DAPI staining. Arrowheads indicate the colocalization of GCaMP6s and anti-NPY staining. **b,** Learning-dependent activity changes of *npy*^+^ dEN axons in the Dc area during Go success, Go failure, and No-go success trials. Note that since fish do not receive punishment during the adaptation session, we referred to these trials as Go goal-reached, Go unreached, and No-go unreached trials. Trial-averaged ΔF/F_0_ patterns (heatmaps) and simultaneous tail movement amplitudes (green lines) from one representative fish are shown. The number of trials used for trial-averaging is indicated above each heatmap, along with the occupancy of each trial type during adaptation, the initial phase of learning, and the after-learning phase. The axonal ROIs are sorted according to their time-averaged response amplitudes during trials in each heatmap. Lightning marks indicate the timing of mild electric stimuli. Other symbols, labels, and annotations are the same as in Fig. 4d. **c,** Learning-dependent changes in ΔF/F_0_ amplitudes of overall dEN axons from multiple fish (mean ± s.e.m.; n = 5 fish). The amplitudes in Go failure trials increased after learning; \**P =* 0.026, two-tailed paired *t*-test. Note that the adaptation session was not included in this analysis due to the absence of electric stimuli, which renders trial outcomes (e.g., Go failure) not equivalent to those in the learning sessions.

**Extended Data Fig. 3:**
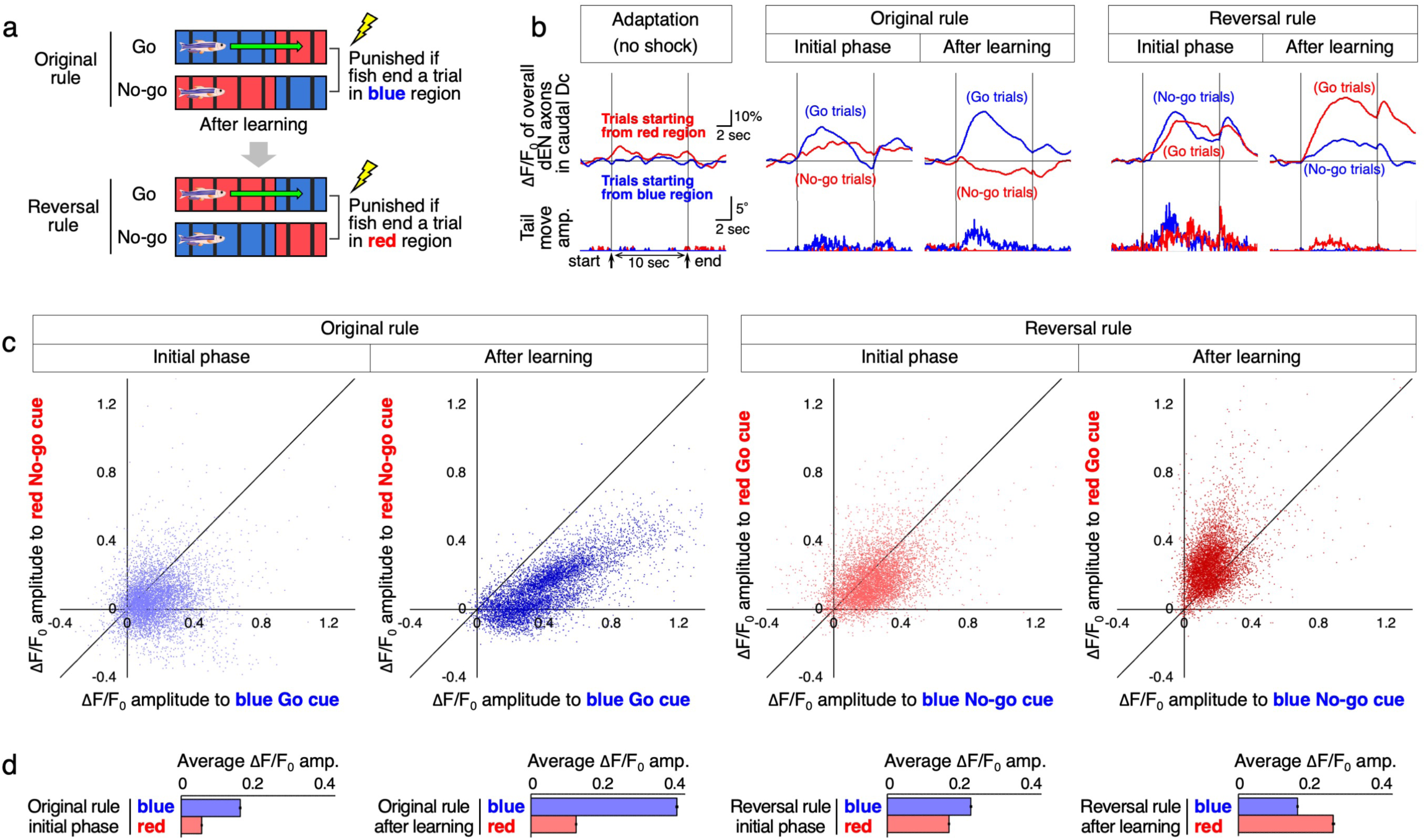
The *npy*^+^ dEN axons’ activity adapts to the rule reversal. **a**, Schematic illustration of the rule reversal experiment. **b,** Activity patterns of overall dEN axons in the Dc area during learning of the original and reversal rules. Trial-averaged ΔF/F_0_ patterns across trials starting from the blue region (blue lines), and across trials starting from the red region (red lines). Data from one representative fish are shown for the adaptation, original rule learning, and reversal rule learning phases. The simultaneous tail movement amplitudes are shown below in the same line colors as the top panels. Horizontal lines indicate ΔF/F_0_ = 0. The dEN axons showed an increased response to the blue cue after learning the original rule. Then, this increased response to the blue cue was diminished, and the dEN axons showed an increased response to the red cue after learning the reversal rule. Scale bars, 10% and 2 sec for ΔF/F_0_; 5° and 2 sec for tail movement. **c,** Learning-dependent changes in distributions of each dEN axon’s ΔF/F_0_ response amplitude to the blue cue against those to the red cue. Each dot indicates an axonal ROI (n = 7414 axonal ROIs from 3 reversal learner fish). Data from the original rule and reversal rule learning phases are shown. **d,** Average ΔF/F_0_ response amplitudes across all the dEN axonal ROIs shown in **c**. Graphs are arranged in the same order as in **c**.

**Extended Data Fig. 4:**
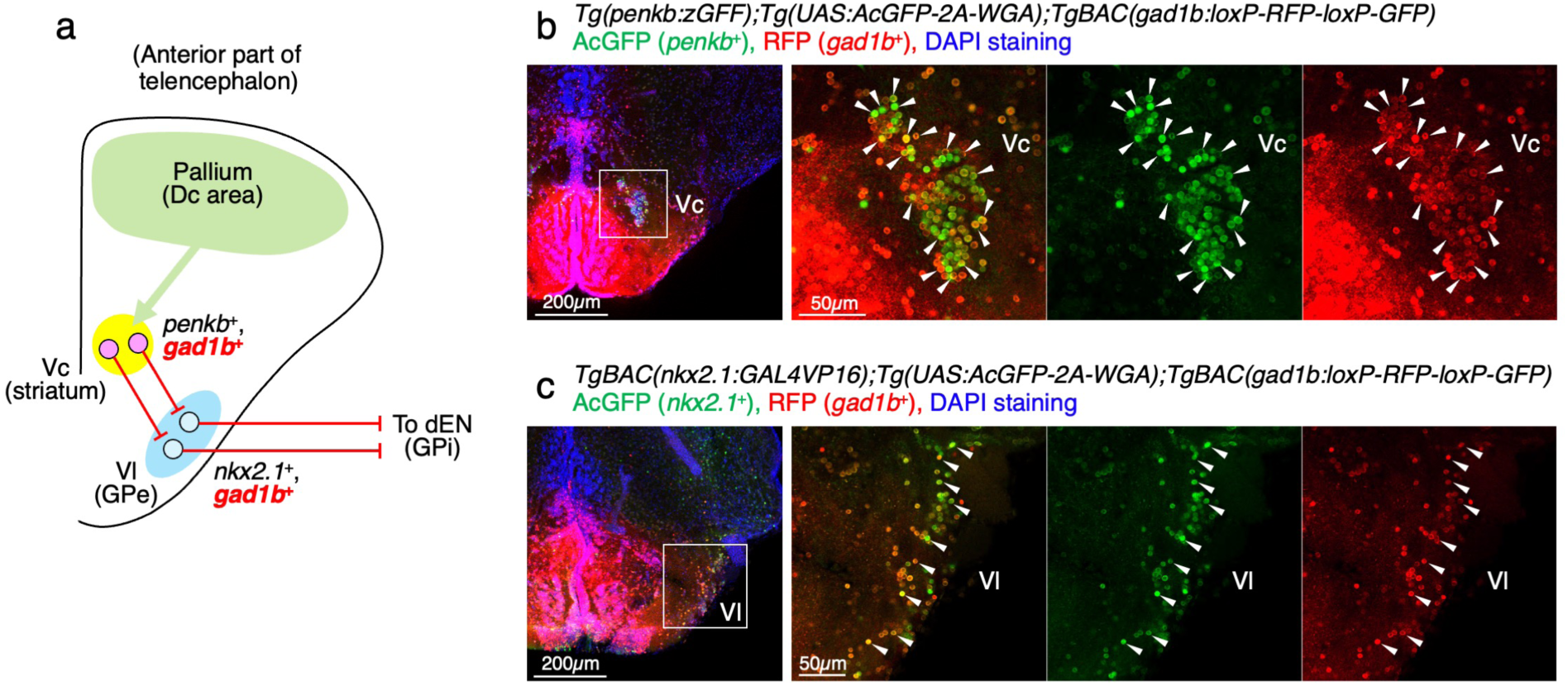
The striatal indirect-pathway neurons are connected to the dEN via two inhibitory projections. **a**, Schematic illustration of the following immunohistochemistry experiments. **b,** Immunohistochemistry for AcGFP (green) expressed in *penkb*^+^ neurons and for RFP (red) expressed in GABAergic neurons in *Tg(penkb:zGFF);Tg(UAS:AcGFP-2A-WGA);TgBAC(gad1b:loxP-RFP-loxP-GFP)* fish. A coronal section at the anterior part of the telencephalon, where the *penkb*^+^ Vc neurons are located,^21^ is shown (see also Fig. 6a). Inset shows the position of the right panels, focusing on the Vc. Arrowheads indicate AcGFP^+^ and RFP^+^ double-positive neurons, indicating that *penkb*^+^ Vc neurons are mainly GABAergic. **c,** Immunohistochemistry for AcGFP (green) expressed in *nkx2.1*^+^ neurons and for RFP (red) expressed in GABAergic neurons in *TgBAC(nkx2.1:GAL4VP16);Tg(UAS:AcGFP-2A-WGA);TgBAC(gad1b:loxP-RFP-loxP-GFP)* fish. A coronal section at the Vl is shown. Inset shows the position of the right panels, focusing on the Vl. Arrowheads indicate AcGFP^+^ and RFP^+^ double-positive neurons, indicating that *nkx2.1*^+^ Vl neurons are mainly GABAergic.

**Extended Data Fig. 5:**
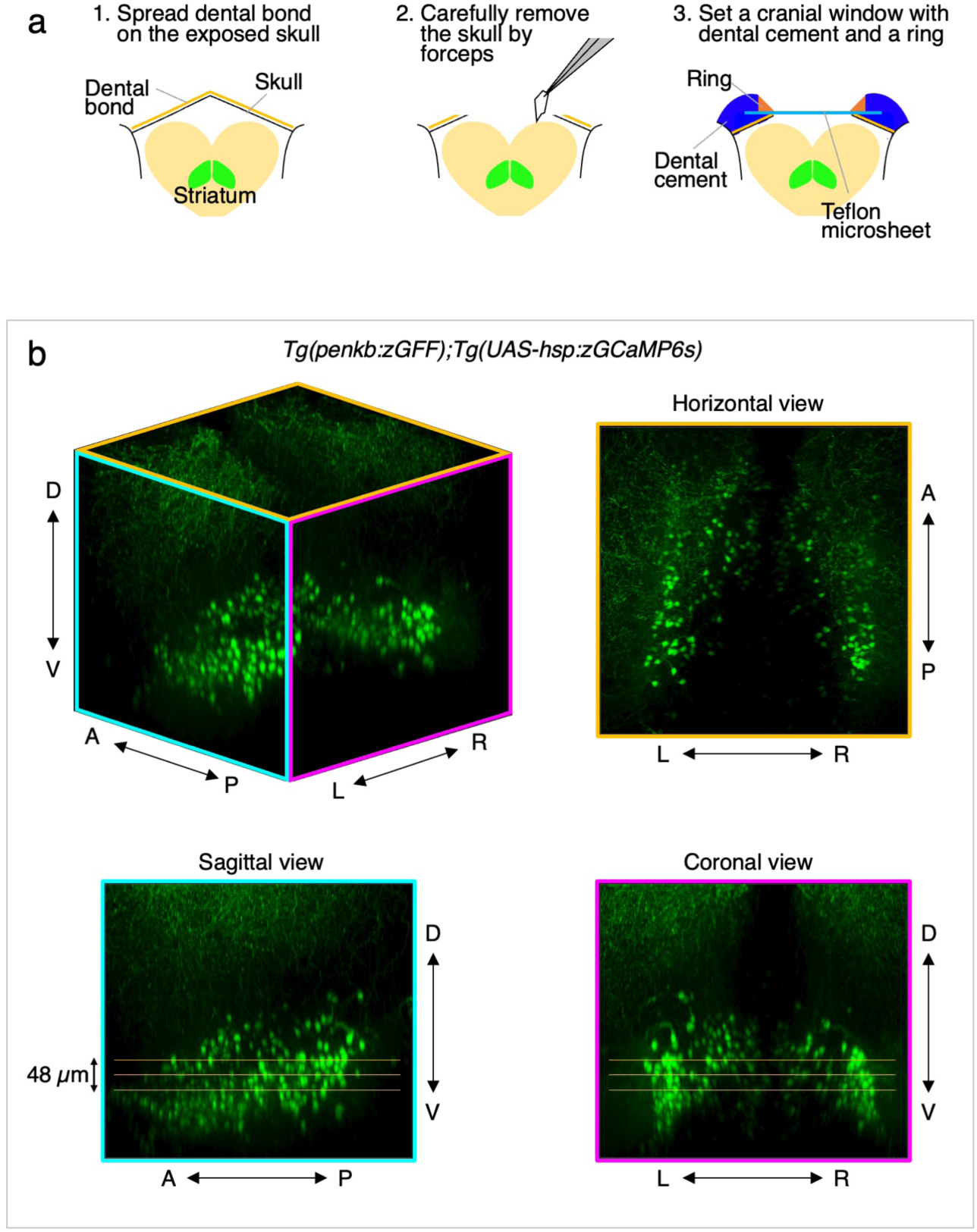
A cranial window protocol using a Teflon microsheet allowed for two-photon imaging of deep brain regions in the adult zebrafish telencephalon. **a**, Illustration of the cranial window surgery using a Teflon microsheet. After head fixation and spreading dental bond on the skull, the telencephalic skull was carefully removed by forceps. Then, a piece of Teflon microsheet and a 3D-printed ring were placed on top of it and carefully embedded with dental cement to cover the surrounding area of the window. See Methods for details. **b,** Two-photon imaging of *penkb*^+^ indirect pathway striatal neurons using the cranial window. The *Tg(penkb:zGFF);Tg(UAS-hsp:zGCaMP6s)* fish were used to evaluate this method. The yellow, cyan, and magenta squares show horizontal, sagittal, and coronal sections of this sample, respectively, which are visualized by maximum intensity projection. The three parallel orange lines indicate typical positions of the three focal planes for the 2-photon calcium imaging of striatal neurons in this study. The intervals between the three focal planes were 24 µm to record activities of different neurons from different focal planes. The top and the bottom planes were 48 µm apart, covering a substantial number of neurons within the view-field. D, dorsal; V, ventral; A, anterior; P, posterior; L, left; R, right.

**Extended Data Fig. 6:**
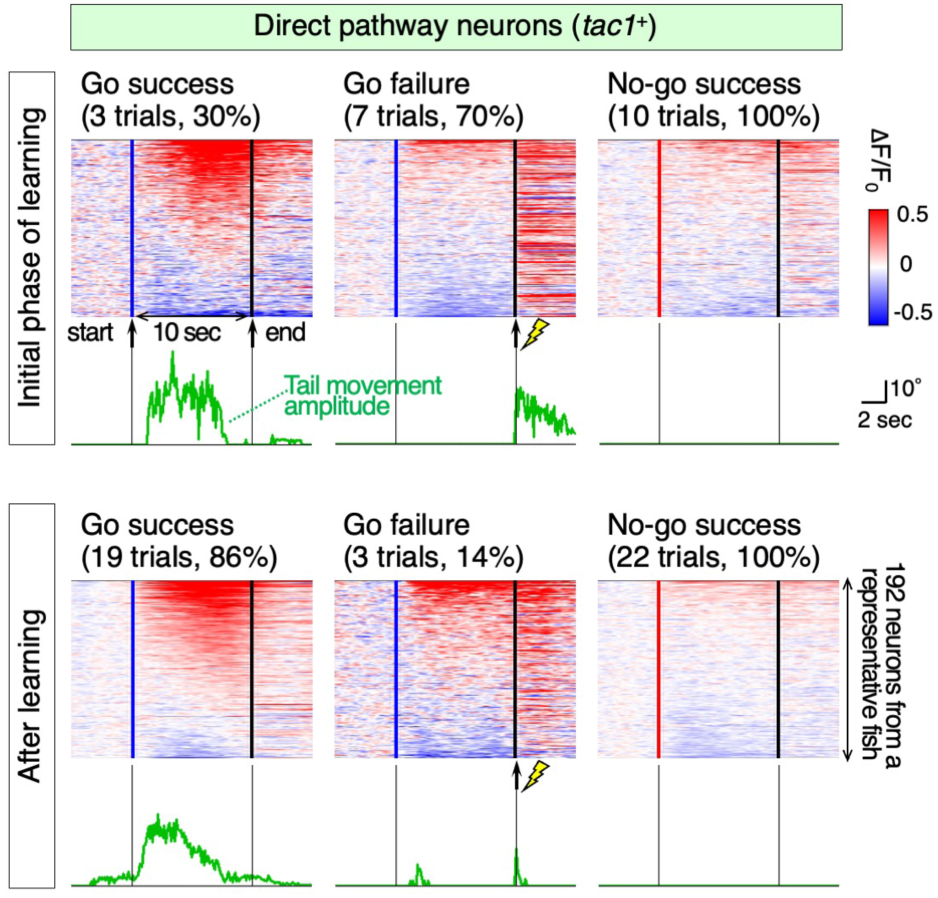
Learning-dependent activity changes of *tac1*^+^ striatal neurons during Go success, Go failure, and No-go success trials. Same as in Fig. 6d, but for *tac1*^+^ direct-pathway neurons. Trial-averaged ΔF/F_0_ patterns (heatmaps) and simultaneous tail movement amplitudes (green lines) from one representative fish are shown. The number of trials used for trial averaging is indicated above each heatmap, along with the Go success, Go failure, and No-go success rates during the learning phase. The neurons are sorted according to their time-averaged response amplitudes during the trials in each heatmap. Blue and red vertical lines indicate the onset of blue and red cues, respectively. Black vertical lines indicate the end of the trial. Lightning marks indicate the timing of mild electric stimuli after failure trials. The results of statistical analysis using multiple fish, including this fish, are shown in Fig. 6f.

**Extended Data Fig. 7:**
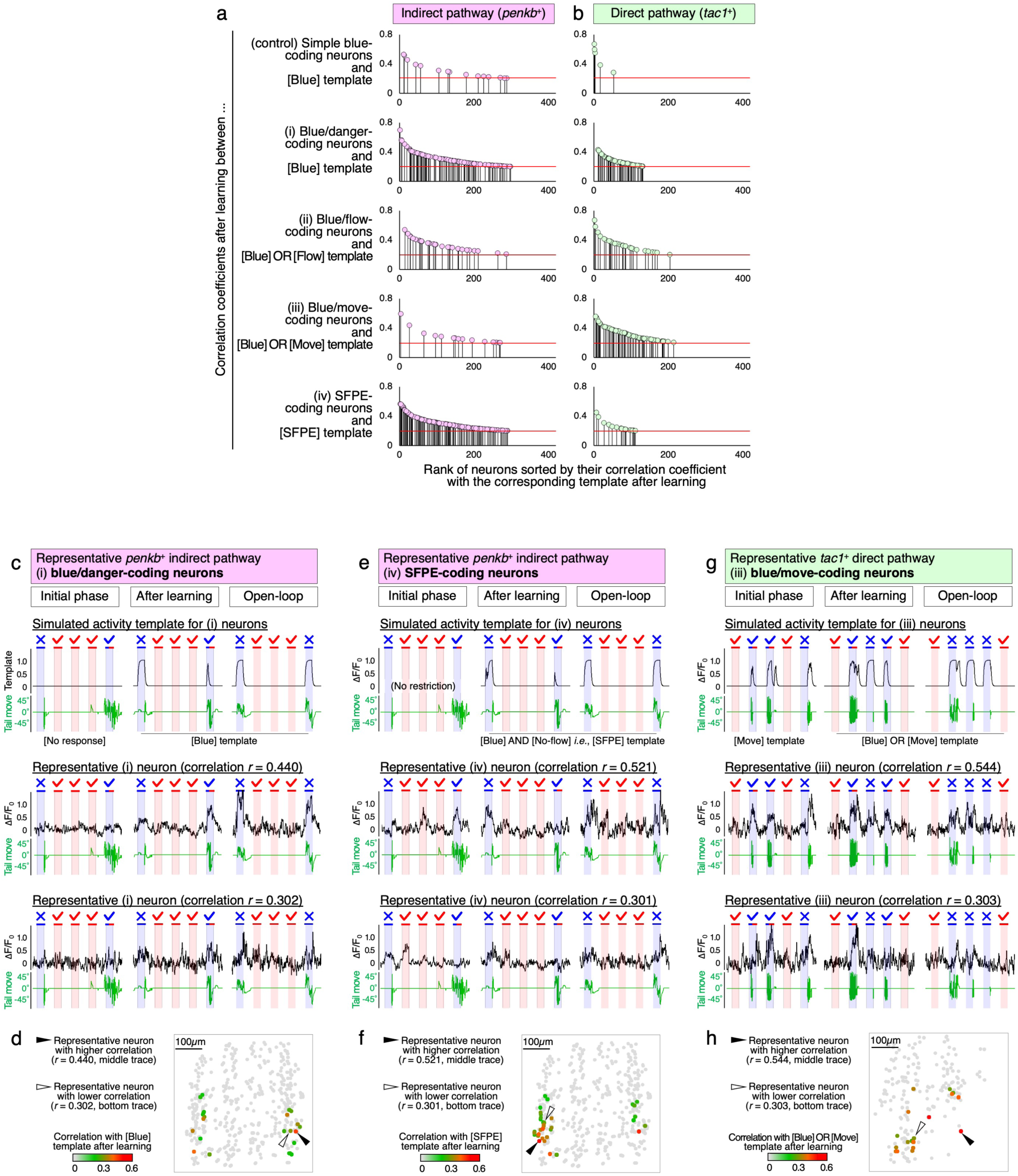
Classification of the striatal neurons that acquired responsiveness to the blue cue after learning. **a**, Distributions of correlation coefficients used for the classification of *penkb*^+^ indirect-pathway neurons. (Top) Correlation coefficients between (control) simple blue-coding neurons and the [Blue] template after learning (see Methods). In each panel, neurons were sorted in descending order by their correlation coefficient with the corresponding template after learning, and the top 400 neurons (out of all 2951 *penkb*^+^ neurons from n = 8 fish) are shown. (Second) Correlation coefficients between (i) blue/danger-coding neurons and the [Blue] template after learning. (Third) Correlation coefficients between (ii) blue/flow-coding neurons and the [Blue] OR [Flow] template after learning. (Fourth) Correlation coefficients between (iii) blue/move-coding neurons and the [Blue] OR [Move] template after learning. (Bottom) Correlation coefficients between (iv) SFPE-coding neurons and the [SFPE] template after learning. Red horizontal lines indicate the classification threshold at 0.2. **b,** Same as in **a**, but for the *tac1*^+^ neurons (1768 *tac1*^+^ neurons from n = 5 fish). **c,** Activity of representative *penkb*^+^ (i) blue/danger-coding neurons with high or low correlation with the corresponding template after learning. (Top) The simulated activity templates for blue/danger-coding neurons. (Middle) The same representative neuron as Fig. 6i, showing high correlation (*r* = 0.440) with the [Blue] template after learning. (Bottom) A representative neuron showing low correlation (*r* = 0.302) with the [Blue] template after learning. For each phase, 5 trials are shown, with the same symbols and annotations as in Fig. 6g. **d,** Distribution of the *penkb*^+^ (i) blue/danger-coding neurons in the representative fish. The same representative fish as in Fig. 6i, right panel. Filled and open arrowheads indicate the positions of the (i) neurons with high and low correlation, respectively. Other symbols and annotations are the same as in Fig. 6i, right panel. **e-f,** Same as in **c-d**, but for representative *penkb*^+^ (iv) SFPE-coding neurons. **g-h,** Same as in **c-d**, but for representative *tac1*^+^ (iii) blue/move-coding neurons.

**Extended Data Fig. 8:**
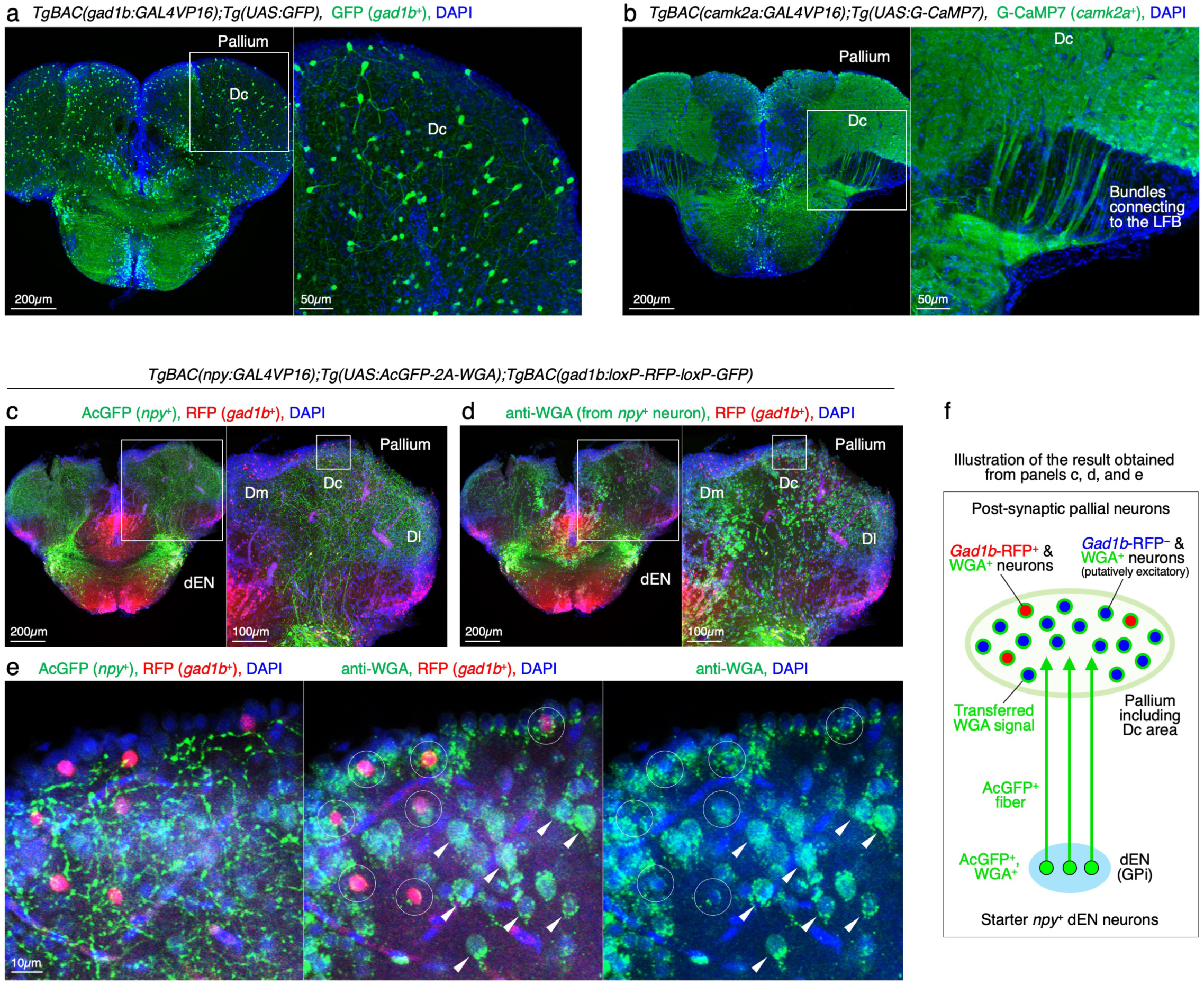
The *npy*^+^ dEN neurons project to most of the Dc neurons, including the inhibitory interneurons. **a**, Immunohistochemistry for GFP (green) expressed in *gad1b*^+^ inhibitory neurons. The inset shows the positions of the right panel. Interneuron-like morphology was observed in the Dc region of the pallium. **b,** Immunohistochemistry for G-CaMP7 (green) expressed in *camk2a*^+^ neurons. This transgenic line was also used for our previous imaging of Dc excitatory neurons in Torigoe et al.^26^ The inset shows the positions of the right panel. Projection-neuron-like staining patterns were observed, which are fiber bundles connecting the Dc region to the lateral forebrain bundle (LFB) in the pallium, in contrast to **a**. **c,** Immunohistochemistry for AcGFP (green) and RFP (red) in *TgBAC(npy:GAL4VP16); Tg(UAS:AcGFP-2A-WGA);TgBAC(gad1b:loxP-RFP-loxP-GFP)* fish. In this line, *npy*^+^ dEN output neurons express AcGFP and wheat germ agglutinin (WGA), a genetically encoded neural tracer,^71^ while *gad1b*^+^ pallial inhibitory neurons express RFP. Large inset shows the position of the right panel. Small inset shows the position of **e**, focusing on the Dc. Dm, medial zone of the dorsal pallium; Dl, lateral zone of the dorsal pallium. **d,** Same as in **c**, but of WGA (green) and RFP (red). The same coronal slice as **c** is shown. WGA is trans-synaptically transferred mainly anterogradely into the postsynaptic cells and, to a lesser degree, retrogradely into the presynaptic cells. Therefore, the projecting fibers of the *npy*^+^ dEN neurons mainly express AcGFP, as shown in **c**, and the neighboring postsynaptic cells accumulate anterogradely transferred WGA, as shown here. **e,** Magnified views of the **c** and **d**. Almost all the Dc neurons accumulated WGA signals, as previously shown in our study.^21^ The RFP^+^ inhibitory Dc neurons (circles) accumulated WGA signals at levels comparable to those observed in the other RFP⁻ Dc neurons (arrowheads), which are putatively excitatory. This result suggests that both inhibitory and excitatory Dc neurons are postsynaptic to the *npy*^+^ dEN output neurons. **f,** Illustration of the results obtained from **c**, **d**, and **e**.

**Extended Data Fig. 9:**
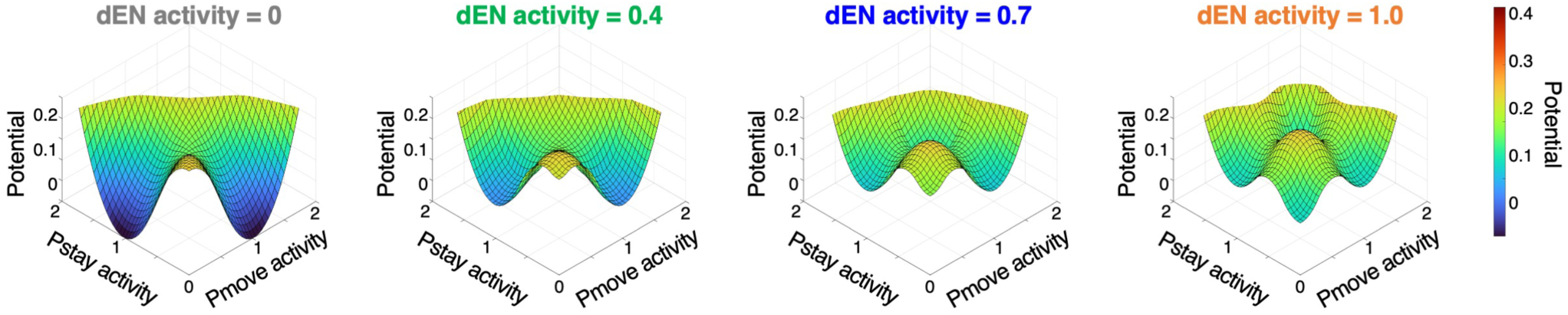
Destabilization dynamics of the Dc activity state by the uniform inhibitory output from the dEN. 3D visualization of the potential landscapes in Fig. 8b, bottom panels. Two-well-like curves appeared when observed from the oblique top view, as we proposed in Fig. 7e-h.

**Extended Data Fig. 10:**
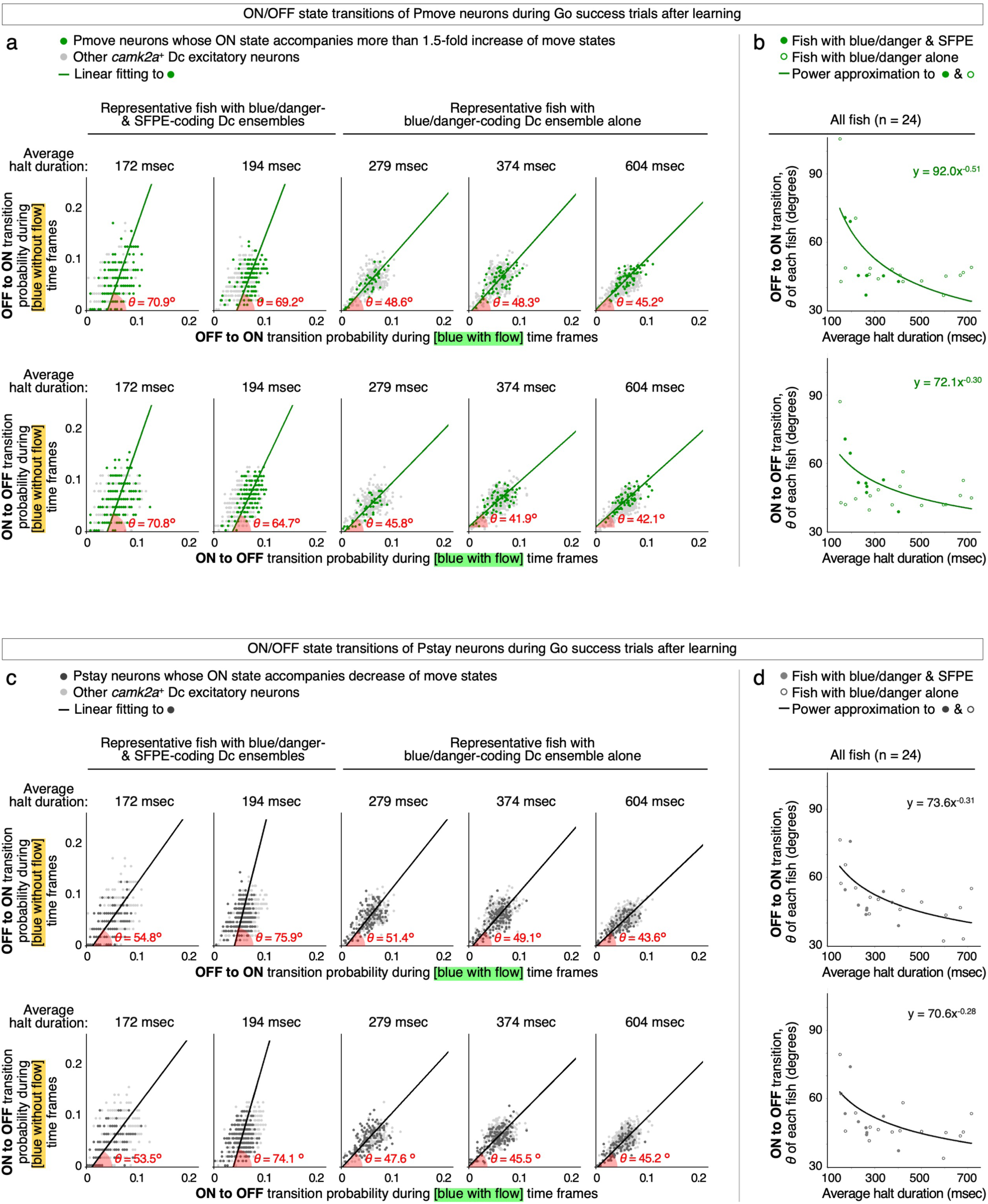
Fish with shorter halt duration show more destabilization of Dc neural states during blue-without-flow time frames. **a**, Analysis of activity ON/OFF state transitions of the *camk2a*^+^ Dc excitatory neurons during Go success trials after learning (original data from Torigoe et al.^26^). (Top) OFF-to-ON transition probabilities for each 100 msec time frame during [blue without flow] time frames were plotted against those during [blue with flow] time frames. Data from 5 representative fish are shown, from the one with the shortest halt duration (leftmost panel) to the one with the longest halt duration (rightmost panel). The first two fish have blue/danger- and SFPE-coding Dc ensembles, and the latter three fish have a blue/danger-coding Dc ensemble alone. Each dot indicates one neuron, and the Pmove neurons are highlighted in green. Green lines are linear fits to the Pmove neurons in each fish. Distribution of transition probability was evaluated as the angle of inclination *θ* of the linear fit; *θ* higher than 45° indicates biased elevation of transition probability in [blue without flow] than in [blue with flow] time frames. (Bottom) Same as in the top panels, but for ON-to-OFF transitions. **b,** Plotting of *θ* from the Pmove neurons against average halt duration in each fish (n = 24 fish), for OFF-to-ON (top) and ON-to-OFF (bottom) transitions. Filled and open circles indicate fish with and without an SFPE-coding Dc ensemble, respectively. Green lines are power approximations across all the fish, and the results of the approximation are shown as the equations (*y* = *ax^b^*). The negative *b* values indicate that the fish with shorter halts tend to have higher *θ*, *i.e.*, biased destabilization during [blue without flow] time frames than during [blue with flow] time frames. **c**-**d,** Same as **a**-**b**, but for Pstay neurons, which are highlighted in dark gray. Similar results were obtained from Pmove and Pstay neurons regardless of OFF-to-ON or ON-to-OFF transitions.

**Supplementary Table 1.** A list of antibodies used in this study.

|  |  |  |  |
|---|---|---|---|
| anti-GFP (chicken polyclonal, 1:3000 dilution) | Aves Labs | GFP-1010, RRID: AB_2307313 | Fig. 4a, 4j, 5b, 6a, 6b, Extended Data Fig. 2a, 4b, 4c, 8a, 8b, 8c-e |
| anti-DsRed (rabbit polyclonal, 1:1000 dilution) | Clontech | #632496 | Fig. 5b, Extended Data Fig. 4b, 4c, 8c-e |
| Anti-Neuropeptide Y (NPY) (rabbit polyclonal, 1:1000 dilution) | SIGMA | #N9528 | Fig. 6a, 6b, Extended Data Fig. 2a |
| Anti-Wheat Germ Agglutinin (WGA) (goat polyclonal, 1:1000 dilution) | Vector Laboratories | #AS-2024 | Extended Data Fig. 8c-e |
| Alexa Fluor® 488 AffiniPure Donkey Anti-Chicken IgY (IgG) (H+L) (1:500 dilution) | Jackson | 703-545-155 | Fig. 4a, 4j, 5b, 6a, 6b, Extended Data Fig. 2a, 4b, 4c, 8a, 8b, 8c-e |
| Alexa Fluor® 647 AffiniPure Donkey Anti-Goat IgG (H+L) (1:500 dilution) | Jackson | 705-605-147 | Extended Data Fig. 8c-e |
| Alexa Fluor® 647 AffiniPure Donkey Anti-Rabbit IgG (H+L) (1:500 dilution) | Jackson | 711-605-152 | Fig. 5b, 6a, 6b |
| Cy™3 AffiniPure Donkey Anti-Rabbit IgG (H+L) (1:500 dilution) | Jackson | 711-165-152 | Extended Data Fig. 2a, 4b, 4c, 8c-e |
| DAPI (1:3000 dilution) | Invitrogen | D1306 | Fig. 4a, 4j, 5b, 6a, 6b, Extended Data Fig. 2a, 4b, 4c, 8a, 8b, 8c-e |

